# Hidden diversity – DNA metabarcoding reveals hyper-diverse benthic invertebrate communities

**DOI:** 10.1101/2022.02.28.481642

**Authors:** Jennifer Erin Gleason, Robert H. Hanner, Karl Cottenie

## Abstract

Freshwater ecosystems, such as streams, are facing increasing pressures from agricultural land use. Aquatic insects and other macroinvertebrates have historically been used as indicators of ecological condition and water quality in freshwater biomonitoring programs; however, many of these protocols use coarse taxonomic resolution (e.g., family) when identifying macroinvertebrates. The use of family-level identification can mask species-level diversity, as well as patterns in community composition in response to environmental variables. Recent literature stresses the importance of robust biomonitoring to detect trends in insect decline globally, though most of these studies are carried out in terrestrial habitats. Here, we incorporate molecular identification (DNA metabarcoding) into a stream biomonitoring sampling design to explore the diversity and variability of aquatic macroinvertebrate communities at small spatial scales. We sampled twenty southern Ontario streams in an agricultural landscape for aquatic macroinvertebrates and, using DNA metabarcoding, revealed incredibly rich benthic communities which were largely comprised of rare taxa detected only once per stream despite multiple biological replicates. In addition to numerous rare taxa, our species pool estimates indicated that after 240 samples from twenty streams, there was a large proportion of taxa present which remained undetected by our sampling regime. When comparing different levels of taxonomic resolution, we observed that using OTUs revealed over ten times more taxa than family-level identification. A single insect family, the Chironomidae, contained over one third of the total number of OTUs detected in our study. Within-stream dissimilarity estimates were consistently high for all taxonomic groups (invertebrate families, invertebrate OTUs, chironomid OTUs), indicating stream communities are very dissimilar at small spatial scales. While we predicted that increased land use would homogenize benthic communities, this was not supported as within-stream dissimilarity was unrelated to land use.

## 1. Introduction

As the effects of climate change become more severe and we enter a sixth mass extinction event (Ceballos et al., 2015), it is now more than ever critical to conserve vulnerable habitats and slow the rate of biodiversity loss. While trends in vertebrate species have been easier to document and traditionally received more attention (e.g., Ceballos et al., 2017, 2020; Rosenberg et al., 2019), the importance of insects and threats towards them have garnered a broader interest in the past decade with the publication of alarming trends in insect decline. For example, Hallmann et al. (2017) estimated that there has been a 75% decline in flying insect biomass in Germany since the late 1980s, and Sánchez-Bayo & Wyckhuys (2019) have predicted that 40% of insect species will be extinct in the next few decades. While there has been debate whether the decline will be as severe as Sánchez-Bayo & Wyckhuys (2019) predicted (e.g., Thomas et al., 2019), it nevertheless remains clear that there is a consistent pattern of insect decline across a broad range of taxonomic groups and habitats in response to climate change and land use (Fourcade et al., 2019; Raven and Wagner, 2021; Wagner et al., 2021). However, it is incredibly challenging to document accurately declines in insect abundance and diversity due to the lack of long-term records (Montgomery et al., 2020) and estimates suggest approximately 86% (upwards of six million) of terrestrial and freshwater eukaryotic species have yet to be described and named by taxonomists (Costello et al., 2013; Mora et al., 2011). As many existing insect population records are limited to a narrow geographic range and lack sufficient temporal scope (e.g., Thomas, 2005), the need for long-term, cost-effective biomonitoring projects to detect changes in insect communities have become increasingly apparent.

Freshwater habitats, such as streams, are particularly threatened by anthropogenic land use and climate change, despite their irreplaceable ecosystem services (Albert et al., 2021; Dudgeon, 2019). The biodiversity of freshwater systems is declining at a faster rate than marine or terrestrial habitats (Reid et al., 2019), stressing how essential stream biomonitoring and conservation projects are to preserve these ecosystems. Biomonitoring assessments often incorporate aquatic invertebrates, which are key bioindicator taxa and sensitive to habitat disturbances (Rosenberg & Resh, 1993). However, these groups present challenges to traditional morphological identification. For example, many immature larval stages cannot be reliably identified to species-level, and routine monitoring is often performed by non-expert taxonomists, which can result in identification errors (Haase et al., 2010, 2006). Due to these constraints, environmental assessments using morphological identifications often use coarse taxonomic resolution (e.g., family-level) as a surrogate for species-level identification, which could potentially mask species-level turnover within a family or not prove sensitive enough to detect impairment (Gleason & Rooney, 2017). The limitations in time and financial resources, large volumes of samples, and either coarse taxonomic resolution or narrow taxonomic focus can be impediments to monitor stream systems exposed to complex physical and chemical stressors.

The above challenges, combined with the need for species detection in an ongoing biodiversity crisis, has prompted research programs which suggest that molecular tools can provide a promising future for freshwater biodiversity assessments (Baird and Hajibabaei, 2012; Blackman et al., 2019; Pauls et al., 2014). Over the past decade, there have been major advancements in molecular identification tools (e.g., high-throughput sequencing or metabarcoding; Cristescu, 2014; Taberlet et al., 2012), which have the potential to be incorporated into biomonitoring programs and ecological research. Metabarcoding has huge potential for environmental assessments as high-throughput sequencing (HTS) platforms can efficiently sequence and identify entire samples (Elbrecht et al., 2017a; Taberlet et al., 2012). In previous studies comparing morphological identification and DNA metabarcoding of invertebrate communities, molecular methods have proven either equally or more effective than traditional approaches at investigating patterns of biodiversity (e.g., Beermann et al., 2018a, 2018b; Elbrecht et al., 2017b; Emilson et al., 2017).

Very speciose taxa, such as the family Chironomidae (non-biting midges, a ubiquitous group of flies with a freshwater larval stage), can especially benefit from metabarcoding applications. Chironomids occupy most freshwater habitats and are often grouped at family-level resolution in assessments (Nicacio and Juen, 2015); however, a mesocosm experiment demonstrated that this family contained 183 operational taxonomic units (OTUs, a proxy for species) when using metabarcoding, and many OTUs had unique associations to water quality and physical habitat parameters (Beermann et al., 2018b).

While metabarcoding provides an avenue for efficient and cost-effective macroinvertebrate identification for biomonitoring programs, the variability of stream habitats can make it challenging to determine whether sampling efforts have been sufficient. Local micro-habitat level conditions, such as riparian vegetation, sediment type, organic matter, and flow regime (e.g., riffle versus pool), are important factors in structuring benthic communities (Beisel et al., 1998; Dobson, 1994). Since stream microhabitats can vary on small spatial scales, this creates a very patchy system in terms of physical stream attributes and resources, and thus affects community composition of aquatic invertebrates (Heino et al., 2004). River and stream microhabitats are unlike typical habitat “patches” due to their branching or dendritic nature (Grant et al., 2007). Isolated habitats, such as ponds, have a clear boundary around the local habitat (from the point of view of aquatic organisms) because it is surrounded by unsuitable terrestrial habitat (Brooks, 2000). Riverine systems offer a unique perspective on community dynamics as there are no discrete boundaries between upstream and downstream habitat patches. The constant flow of water can allow for continuous dispersal, and defining a “local” community in streams thus poses a challenge for biomonitoring programs as it is unclear how much sampling effort is necessary to represent accurately diversity within a site. This spatial extent knowledge gap, combined with the potentially unknown taxonomic diversity of benthic invertebrates, can be uniquely answered by sampling methods based on metabarcoding principles to inform biomonitoring protocols and efficiently record biodiversity in stream systems.

Small-scale variations in stream habitats can be caused by both natural and anthropogenic processes, although loss of microhabitats (e.g., habitat homogeneity) is often associated with adjacent agricultural land use, which results in channelization and reduction of riparian vegetation (Allan, 2004). Heterogeneous habitats tend to support higher species richness (Ortega et al., 2018; Stein et al., 2014), and it is perhaps unsurprising that increasing agricultural land use in the surrounding catchment area can cause taxonomic and functional homogenization in aquatic macroinvertebrates, resulting in more similar communities of ‘tolerant’ taxa (Siqueira et al., 2015, Martins et al., 2017). However, the relationship between beta diversity (a measurement of community dissimilarity) and both land use and habitat heterogeneity in stream macroinvertebrates is unclear and often case dependent. In Finland, Heino et al. (2013) determined that heterogeneity was not a significant predicator of invertebrate beta diversity in streams using a mix of species and genus-level identifications. Additionally, research using predominately genus-level identifications by Petsch et al. (2021) concluded that land use did not cause homogenization of stream invertebrates in boreal (Finland) or subtropical (Brazil) regions. However, contrasting patterns have been detected in New Zealand, where habitat heterogeneity was a strong driver of beta diversity in stream invertebrate communities (Astorga et al., 2014) and in North America (Maryland, USA) where Maloney et al. (2011) detected a negative relationship between beta diversity and increased pasture and crop cover.

To address the spatial knowledge gap in streams, field collection needs to occur at an appropriate scale. The above studies of stream invertebrate beta diversity patterns are performed at large spatial scales (e.g., comparing beta diversity between major watersheds or geographic regions), and there are few stream studies that explore spatial resolution at small scales (e.g., microhabitat level, but see Costa & Melo, 2008 and Heino et al., 2013). In addition to this, many of these studies are performed at variable taxonomic resolution using morphological identification (a mix of family, genus, and rarely species), focus on specific taxonomic groups (e.g., Heteroptera only; Dias-Silva et al., 2020), or exclude diverse insect groups such as chironomids (e.g, Astorga et al., 2014; Costa & Melo, 2008; Petsch et al., 2021). Next-generation sequencing methods have been extremely effective at revealing incredibly diverse communities of terrestrial invertebrates (D’Souza and Hebert, 2018; Maggia et al., 2021; Steinke et al., 2021a, 2021b), and it is likely that using molecular identification to explore similar patterns in streams will provide greater insight into both alpha and beta diversity patterns.

In this study, we used DNA metabarcoding to determine how variable benthic invertebrate communities are at small spatial scales across three time points in a single year. We determined the importance of taxonomic resolution in revealing biodiversity patterns by performing all analyses at both family-level and OTU-level resolution. As chironomid OTUs generally comprise high levels of diversity in freshwater samples, we also repeated all analyses using only OTUs from this family. We assessed whether overall taxonomic richness is linked to land use and calculated within-stream dissimilarity to determine if small-scale changes in community composition are influenced by agricultural activity. We hypothesized that agricultural landscapes homogenize stream communities and will result in more uniform benthic communities due the loss of habitat complexity, and therefore predicted that within-stream dissimilarity will decrease (e.g., more homogenous communities) as the percentage of agricultural land use in the catchment increases. We also explored how stream biodiversity estimates change between different levels of taxonomic resolution by calculating rarefaction curves and estimating the total regional species pool, calculating the sampling coverage at each stream site and determining the percentage of a local community made up by rare taxa in order to inform future sampling efforts.

## 2. Methods

### 2.1 Site selection and stream sampling

We collected benthic macroinvertebrates from twenty streams in southern Ontario across three time periods (May, July, and September 2019; Figure 1). We selected streams on a continuum of surrounding land use, and sites were located either on Conservation Authority property or privately owned land (farm sites), and additionally were required to be wadable and wet for the entire study period. We used the Ontario Flow Assessment Tool (OFAT; Government of Ontario, 2020) to determine stream watershed boundaries in ArcGIS v. 10.6.1 (Esri, 2020) and the Ontario Land Cover Compilation v. 2.0 (Ontario Ministry of Natural Resources and Forestry, 2016) to determine the percentage of agriculture land use (cropping) surrounding each stream site.

**Figure 1.**
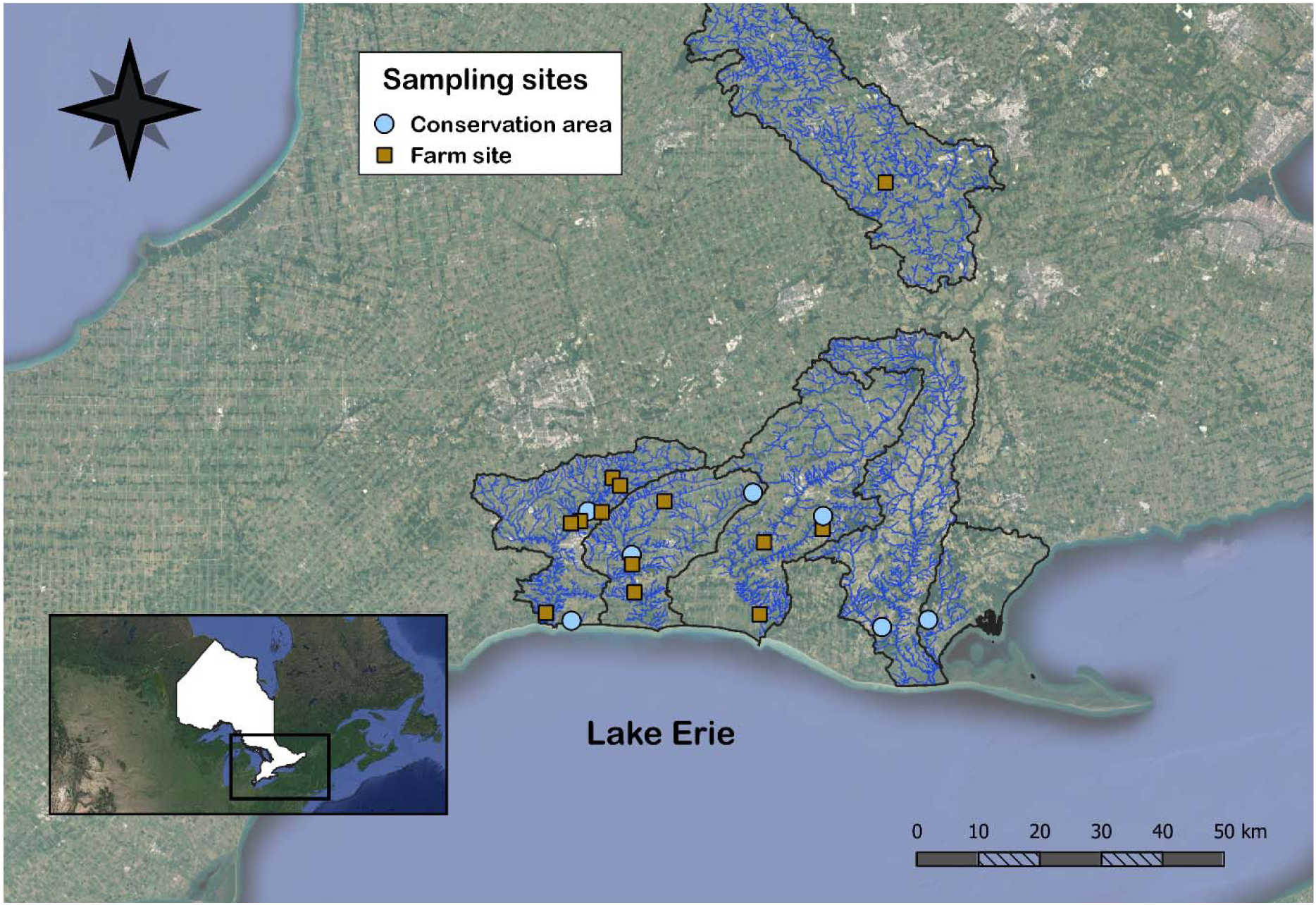
A map of stream sampling sites in conservation areas (blue circles) and privately owned farms (orange squares) in southern Ontario, Canada, along the north shore of Lake Erie. The map was created using QGIS (Q GIS DevelopmentTeam, 2020). The black outlines demonstrate quaternary watershed boundaries where we sampled. The inset map shows the province of Ontario in white with our study region outlined in a black rectangle.

For field collection of aquatic macroinvertebrates, we collected four biological replicates within each stream (i.e., four bulk samples per stream) by selecting four transects that were approximately 10-20 meters apart and positioned downstream to upstream to avoid contamination from sampling-related disturbance. We placed transects to include multiple microhabitats if present (e.g., riffles and pools, different substrates) and collected benthic macroinvertebrates and associated habitat information based on the Ontario Benthic Biomonitoring Network (OBBN; Jones et al., 2007) and the Ontario Stream Assessment Protocol (OSAP; Stanfield, 2017). Each sample consisted of a 3-minute travelling kick-and-sweep using a 500 µm D-net across the width of the stream. We then transferred the bulk sample to a 500 µm mesh sieve for rinsing and the removal of large debris, before storing in a sample container and preserving in 95% ethanol on site. We kept the invertebrate samples in a chilled cooler until transfer to the lab on the same day, where they were stored in a 4°C fridge until further processing. All of our sampling equipment (e.g., nets, sieves, forceps, waders) were cleaned with a 10% bleach solution and rinsed with de-ionized water (DI) between sites. In total, we collected four biological replicates per stream and 80 samples each month, for a total of 240 bulk samples.

### 2.2 Sample sorting and DNA extraction

Bulk macroinvertebrate samples were rinsed with DI water over a sterilized 500 µm sieve and sorted under a dissection microscope. Benthic macroinvertebrates were removed from sample debris and placed in a sterile 20 mL tube containing 95% ethanol and ten 4 mm diameter steel beads. As many bulk samples were very large, we used a subsampling approach based on equal effort by stopping sorting after 4 hours had elapsed. The unsorted portion of sample was placed on a white grid and scanned for two minutes for any rare taxa that had not been encountered during the initial subsampling. After the specimens had been removed from debris, they were air dried for one week while covered with a kimwipe to prevent any infall. The dry biomass of each sample was recorded, and then samples were homogenized using an IKA Tube Mill (IKA, Staufen, Germany) at 4000 rpm for 15 minutes. Smaller samples were ground in a 2 mL sterile tube with two steel beads using a TissueLyser II (Qiagen, Hilden, Germany) at 30 hz for one minute. We subsampled 20 mg (± 1 mg) of ground tissue into a sterile 2 mL tube and used a DNeasy Blood & Tissue Kit (Qiagen, Hilden, Germany) following manufactures’ guidelines for DNA extraction, follow by quantification using a Qubit 3.0 Fluorometer (ThermoFisher Scientific, MA, USA). Several samples contained less than 20 mg of tissue, and the entire sample was used for DNA extraction in place of sub-sampling.

### 2.3 PCR amplification and library preparation

We selected the mitochondrial cytochrome c oxidase subunit I (CO1) gene as our marker and targeted a 421 base pair region to amplify in our initial PCR reaction. We used the Qiagen multiplex PCR kit (Qiagen, Hilden, Germany) as our master mix and selected a primer pair (BF2 + BR2; Elbrecht and Leese, 2017) that has been successful at amplifying a broad range of invertebrate taxa, including aquatic invertebrates collected from our study region (Gleason et al., 2020; Persaud et al., 2021). Each reaction consisted of 2.5 µL DNA extract, 12.5 µL of 2x Qiagen master mix, 9 µL of molecular water, and 0.5 µL of each primer (BF2 + BR2, initial concentration of 0.2 µM) for a total reaction volume of 25 µL. Our thermocycling profile followed Qiagen’s manufacturer’s protocol: a 95°C initial denaturation for fifteen minutes, followed by 25 cycles of 94°C for 30 seconds, 50°C for 90 seconds, 72°C for 60 seconds, and a final extension at 72°C for ten minutes and visualized using precast 2.0% agarose e-gels (E-Gel 96 SYBR Safe DNA stain; ThermoFisher Scientific, MA, USA). Each sampling period (e.g., month) consisting of 80 samples were prepared in their own plate, in addition to 6 PCR negative controls, 1 sequencing negative control, and 1 extraction negative control. We included 8 PCR technical replicates per plate to ensure PCR reproducibility and explicitly selected samples of both lower and higher invertebrate abundance. This resulted in three 96-well PCR plates consisting of 240 samples, 24 technical replicates, and 24 negative controls. The resulting PCR products were purified using NucleoMag NGS clean up and size select magnetic beads (Macherey-Nagel, USA) with an 0.8x ratio of beads to PCR product as per Milián-García et al. (2021).

A second indexing PCR reaction was prepared using Illumina indexing primers (Set A) to tag samples for library preparation. Here, we prepared a 50 µL PCR reaction using 5 µL of our purified PCR product, 25 µL of 2x Qiagen master mix, 10 µL of molecular water, and 5 µL each of forward and reverse indexing primers (initial concentration 10 µM) based on Illumina’s standard indexing protocol. The thermocycling profile included an initial denaturation at 95°C for fifteen minutes, 8 cycles of 95°C for 30 seconds, 55°C for 30 seconds, 72°C for 30 seconds, and a final extension of 72°C for five minutes. After e-gel visualization to confirm amplification, we again purified the PCR products using NucleoMag beads (0.6x ratio, Milián-García et al., 2021). After a final visualization, we submitted the prepared libraries to the Advanced Analysis Center at the University of Guelph. Each plate was normalized, pooled, and sequenced separately on the Illumina MiSeq platform for a total of three separate runs.

In some cases, samples did not perform well in sequencing and were subsequently filtered out of the dataset based on low sequence read (36 samples filtered out with fewer than 80k sequences). Most of the failed samples came from the same streams and had lower-than-average DNA concentration, and we re-ran these samples following the same protocol as above, but instead increased the template volume to 5 µL in the initial amplification PCR and added an additional 10 cycles to the thermocycling program and submitted a fourth plate for sequencing as above to replace failed samples.

### 2.4 Bioinformatics pipeline

We used the bioinformatics platform JAMP v. 0.67 (http://github.com/VascoElbrecht/JAMP) to process the raw sequence data. The protocol is listed in detailed in Persaud et al. (2021), but in brief this involved paired-end merging of de-multiplexed reads using USEARCH (v. 11.0.6668; Edgar, 2010) followed by trimming primer sequences from reads using cutadapt (v. 1.15; Martin, 2011.). We assessed sequence size by filtering out any that were more than ten base pairs longer or shorter than our target (421 bp) and filtered out low-quality sequence with expected errors ≥ 1. We used USEARCH (v. 11.0.6668; Edgar, 2010) to cluster sequences into Operational Taxonomic Units (OTUs) using a 97% similarity threshold, and OTUs with less than 0.01% abundance across all samples were filtered out (e.g., Beermann et al., 2018b; Steinke et al., 2021a). We matched our OTUs to the Barcode of Life Database reference sequence library (BOLD; Ratnasingham and Hebert, 2007) using the Python program BOLDigger (Buchner and Leese, 2020). Raw sequences are available on NCBI SRA (Accession: PRJNA783201) and our final OTU table with sequence reads per sample and associated taxonomic metadata are available as supplementary information.

### 2.5 Data quality control

All of our statistical analyses and figures were performed using R version 4.0.3 (R Core Team, 2020), and all plots were created using the package ggplot2 (v.3.3.3; Wickham, 2020). We used the R package metabaR (v. 1.0.0; Zinger et al., 2021) to assess the quality of our metabarcoding data, including confirming sequencing depth was appropriate and checking for contamination on sequences present in our negative controls (see Appendix 1 for further details). A sequence was identified as a contaminate if it had a relative abundance that was highest in a negative control (as no other DNA should be present in negative controls, contaminants should be preferentially amplified). If more than 10% of reads within a sample corresponded to contaminant sequences, the sample was removed from the dataset. Based on the small number of reads in sequencing controls, we tested multiple filtering thresholds to lower the influence of tag jumps and prevent false positives. The abundance of an OTU in sample was changed to zero if the relative abundance of that OTU was less than 0.001% of the total abundance of that OTU in the entire dataset. We then filtered out samples with low sequence reads (less than 1 SD below average sequence read; 87344) and assessed the quality of technical replicates for reproducibility based on Bray-Curtis distances within and between samples (e.g., contrasting the dissimilarities in OTU composition). A sample was flagged as a failed if the distance within a sample (e.g., between technical replicates) was greater than the threshold of the intersection value for within and between sample distances. We filtered out any non-target taxa and retained only arthropods, annelids and molluscs which were the three must abundant phyla in terms of both total sequence reads and OTU counts. We did not include nematodes in any further analysis as we did not obtain many sequences or OTUs matching to this group, likely due to a combination of the small size of some species and primer bias. Finally, we filtered out poor-quality taxonomic matches (< 90% match to reference database). After cleaning the data using metabaR, we calculated the total number of sequences and OTUs that were removed from the dataset during this process. See Appendix 1 for additional details and figures for the above protocol.

### 2.6 Statistical analysis

To assess the influence of taxonomic resolution on ecological patterns, we analyzed the data first at OTU-level and then repeated all analysis with data scaled back to family-level resolution. We also repeated all analyses using only the chironomid OTUs, the most abundant and species-rich family in our dataset. All data were analyzed at all three time points in our data set (May, July, September) and at three taxonomic levels (all invertebrate OTUs, all invertebrate families, chironomid OTUs only).

We first used a two-way ANOVA to determine if there were significant differences in taxon richness between stream type (e.g., located in a conservation area or on private property) and between sampling months. To assess how within-stream dissimilarity is influenced by surrounding land use, we calculated the Raup-Crick index between the four biological replicates (transects) within a strean using the ‘raupcrick’ function with 999 simulations in the R package vegan (v. 2.5-7; Oksanen et al., 2016) and then took the mean of all pairwise comparisons as the dissimilarity value for that site. Using Raup-Crick as a dissimilarity index is ideal for metabarcoding data as it treats the number of sequence reads per OTU as binary (e.g., presence/absence) as opposed to an abundance value, and it is based on occurrence probabilities in proportion to frequency of a species occurrence and thus should be robust to the influence of rare taxa and large differences in richness between sites. We then used linear regressions to determine if there was a significant effect of the percentage of agricultural land use surrounding the stream on the overall dissimilarity.

We used the R package iNEXT (v. 2.0.20; Hsieh et al., 2020) to generate species rarefaction and extrapolation curves to estimate the total landscape-scale diversity in each sampling period (month) for each taxonomic resolution group. To estimate how many undetected taxa remained at each stream site (e.g., locally), we calculated the number of expected total taxa at each based on Chao’s equation using the ‘specpool’ function in the R package vegan (Oksanen et al., 2017) and plotted this as the percentage of coverage achieved by dividing the observed number of taxa over the expected number of taxa. Finally, to assess how many rare taxa make up a local community, we calculated the percentage of taxa which only occurred in one (of four) biological replicates at each stream site. We performed two-way ANOVAs on both datasets (the percentage of coverage and the percentage of rare taxa) to determine if there were significant differences between sampling months or levels of identification. To determine which variables differed significantly, we used Tukey’s multiple comparison of means to calculate adjusted p-values using the ‘TukeyHSD’ function.

## 3. Results

### 3.1 Summary and taxonomic richness

We received 74,771,159 sequence reads from the four MiSeq runs (average of 18,893,999 sequences per run), and after post-bioinformatic processing our samples contained a total of 51, 334, 969 reads (average per sample 161,430 ± 69,286) and 2276 OTUs (average 72 per sample ± 33), while all negative controls combined had only 1954 reads (average per control 57 ± 16) and 241 OTUs (average per control 16 ± 13). Notably, our OTU table originally contained 5597 OTUs which were filtered out after clustering for not meeting the 0.01% abundance threshold (these OTUs corresponded to only 33,840 sequences in total). Three OTU sequences were flagged as contaminants through metabaR (Zinger et al., 2019), two of which were unidentified algae and one matched to a species of maple tree (*Acer* sp.), indicating that the most likely cause of the small amount of sequences in the negative controls were tag jumps during sequencing. No samples were flagged as contaminated, and all technical replicates passed our reproducibility criteria. Three samples were removed from the dataset for sequencing depth lower than one standard deviation of the mean (< 86k sequences). The most common reason for an OTU to be removed from the final dataset was a match of less than 90% to the sequence reference database (BOLD). Post metabaR (Zinger et al., 2021), our cleaned dataset contained a total of 49,072,214 high quality reads (177,156 per sample ± 47,828.58) and 1663 macroinvertebrate OTUs (61 per sample ± 29). We created two more data frames from the cleaned data, one with OTUs scaled back to family-level resolution (149 families total) and one containing only OTUs from the family Chironomidae (560 OTUs), as chironomids contained a third of the total diversity within the invertebrate dataset.

There was no significant influence of site type (CA versus farm; F_1,56_ = 0.67, *p* = 0.4) or sampling month (F_2, 56_ = 0.74, *p* = 0.5) on average OTU richness (Figure 2). Likewise, there was no effect of either site type or sampling month on average family richness (type: F_1,56_ = 0.16, *p* = 0.69; month: F_2,56_ = 1.31, *p* = 0.28) or average chironomid OTU richness (type: F_1,56_ = 0.13, *p* = 0.72; month: F_2,56_ = 1.66, *p* = 0.20; Figure 2).

**Figure 2.**
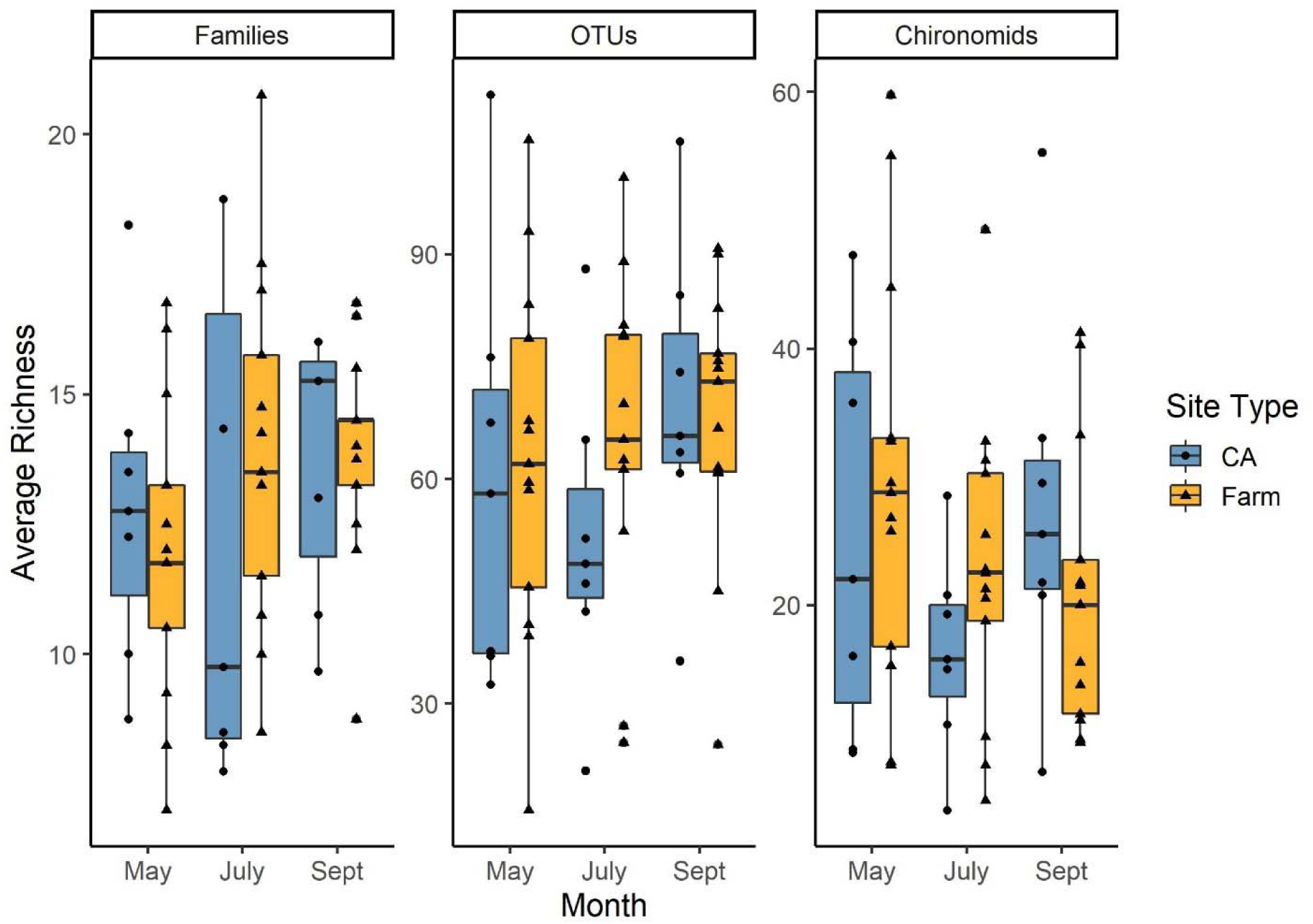
The average taxonomic richness per site in each category (conservation area - CA), n = 7, or farm, n = 13) over the sampling period (May, July, Sept, total n = 237). There was no significant effect of site type or sampling month on taxonomic richness for any group. Note the different Y-axis scales in the 3 figures.

### 3.2 Site dissimilarity and agricultural land use

Within-site Raup-Crick dissimilarity was not significantly correlated with the percentage of agricultural land use for any month or level of identification (all families, all OTUs, chironomid OTUs; Figure 3). Mean dissimilarity was generally consistent across sampling months for family resolution but was lowest in May for the OTU-level datasets, and dissimilarity was generally high for all sites (i.e, most values within range 0.5-1.0; Figure 3).

**Figure 3.**
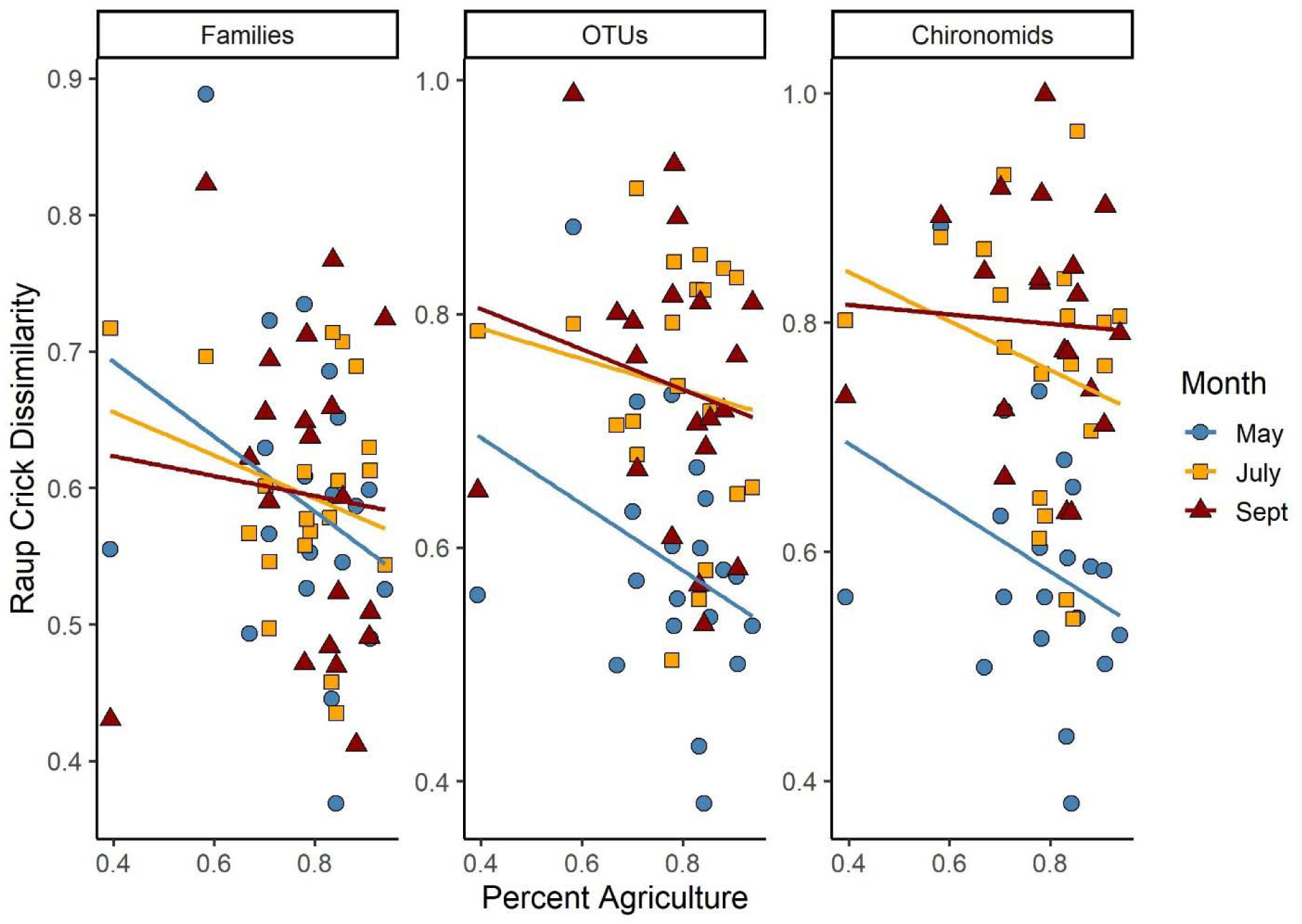
The mean Raup-Crick index of all pairwise comparisons within a site is plotted against the percentage of agricultural land use surrounding the stream site. There was no significant relationship between dissimilarity and agriculture for any identification level or sampling month.

### 3.3 Rare and undetected taxa

We collected a total of 80 samples for each sampling period, but the species rarefaction curves did not level off (Figure 4). Extrapolations estimated that at 150 sampling units, the number of new taxa would begin to level off for family-level identification and for chironomid OTUs. Based on Chao’s equation, we determined that each sampling month collected approximately 71% of total invertebrate families present, 69% of all invertebrate OTUs present, and 77% of chironomid OTUs present, and at the stream level this value ranged dramatically from 29-94% for families, 27-85% for all invertebrate OTUs, and 14-90% for chironomid OTUs, indicating that many taxa can be missed locally (Figure 5). We found that there were significantly more ‘undetected’ taxa in the OTU datasets compared to the family-level dataset (F_2,175_ = 5.65, *p* = 0.004) and observed that sampling month had a significant effect on the percentage of coverage (F_2,175_ = 4.25, *p* = 0.01), with September being significantly different from May and July (adjusted *p =* 0.02).

**Figure 4.**
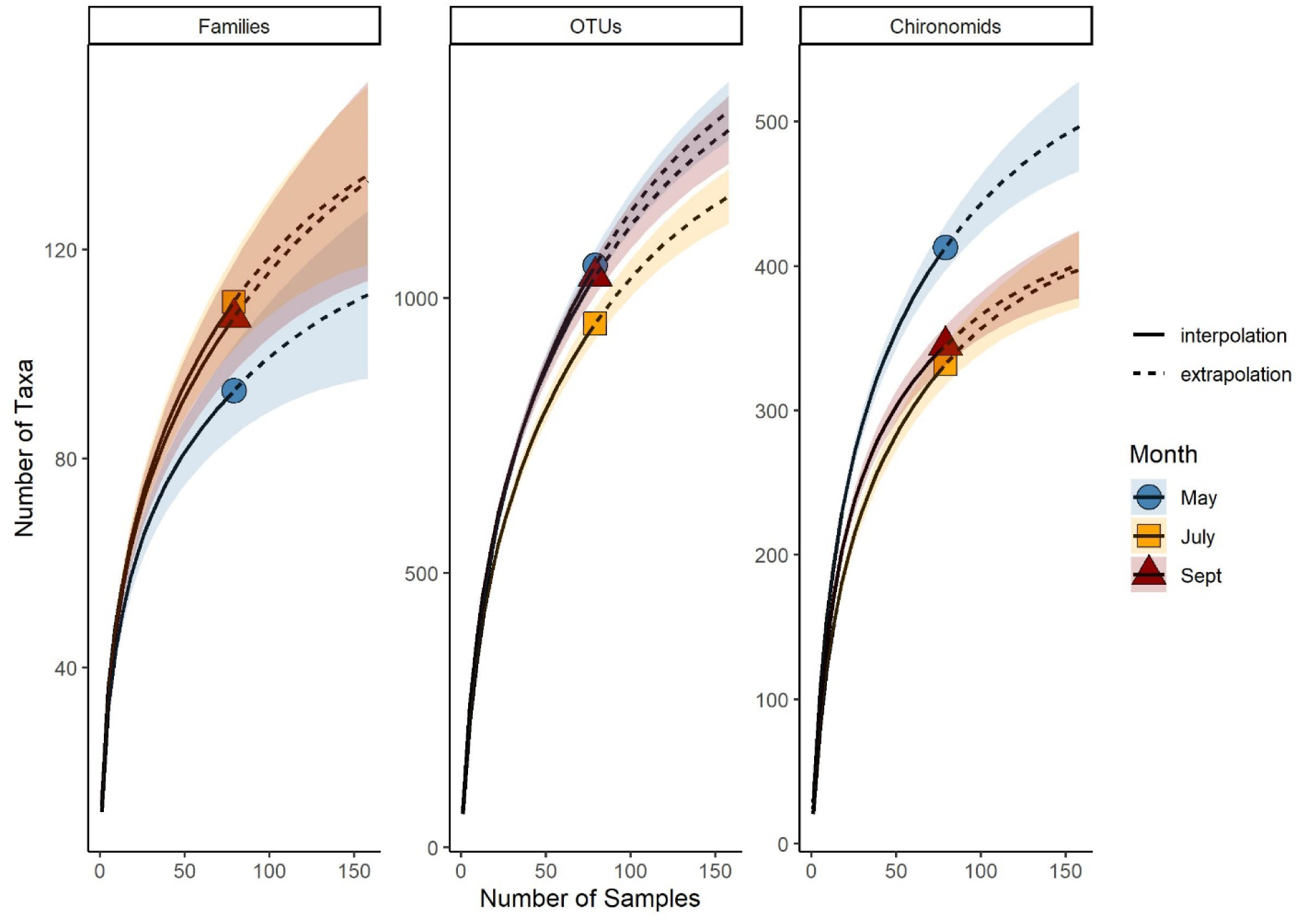
Rarefaction curves for the number of taxa collected each sampling month where the solid line represents interpolated richness from our samples and the dashed line is an extrapolation based on expected number of taxa with continued sampling. Symbols indicate the point where our sampling stopped (n = 80). Note difference in y-axis scales.

**Figure 5.**
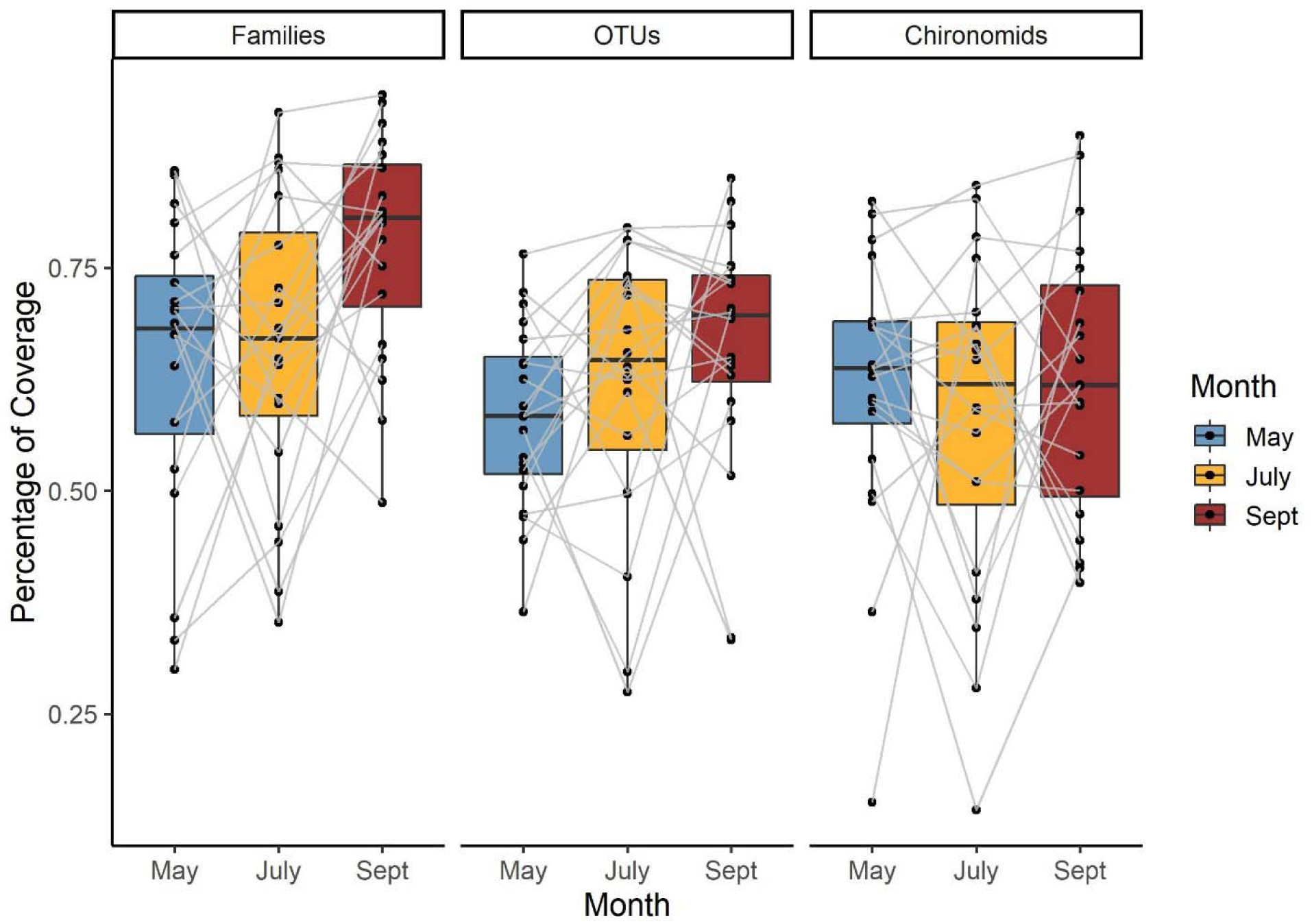
Estimated percentage of taxonomic coverage achieved at each stream by dividing the observed number of taxa over the extrapolated number of taxa expected based on Chao’s equation for each sampling month. Grey lines connect the same stream over time. There were significantly more ‘undetected’ taxa in the OTU datasets compared to the family-level dataset (F_2,175_ = 5.65, *p* = 0.004). Sampling month also had a significant effect on the percentage of coverage (F_2,175_ = 4.25, *p* = 0.01), with September having the least percentage of undetected taxa.

In addition to undetected taxa, we also calculated the proportion of rare taxa at each stream to determine how variable streams are at small spatial scales. Streams had on average 59% (range 40-87%) of invertebrate OTUs that were only detected in one of four biological replicates, indicating a high degree of turnover within a single site (Figure 6). Likewise, there was an average of 46% of unique taxa per stream (range 24-71%) for invertebrate families and 61% (range 31-94%) for chironomid OTUs (Figure 6). We observed significantly more rare taxa in the two OTU datasets compared to family-level resolution data (F_2,175_ = 26.1, *p* < 0.001), and found no significant effect of sampling month on the percentage of rare taxa locally (F_2,175_ = 2.09, *p* = 0.127).

**Figure 6.**
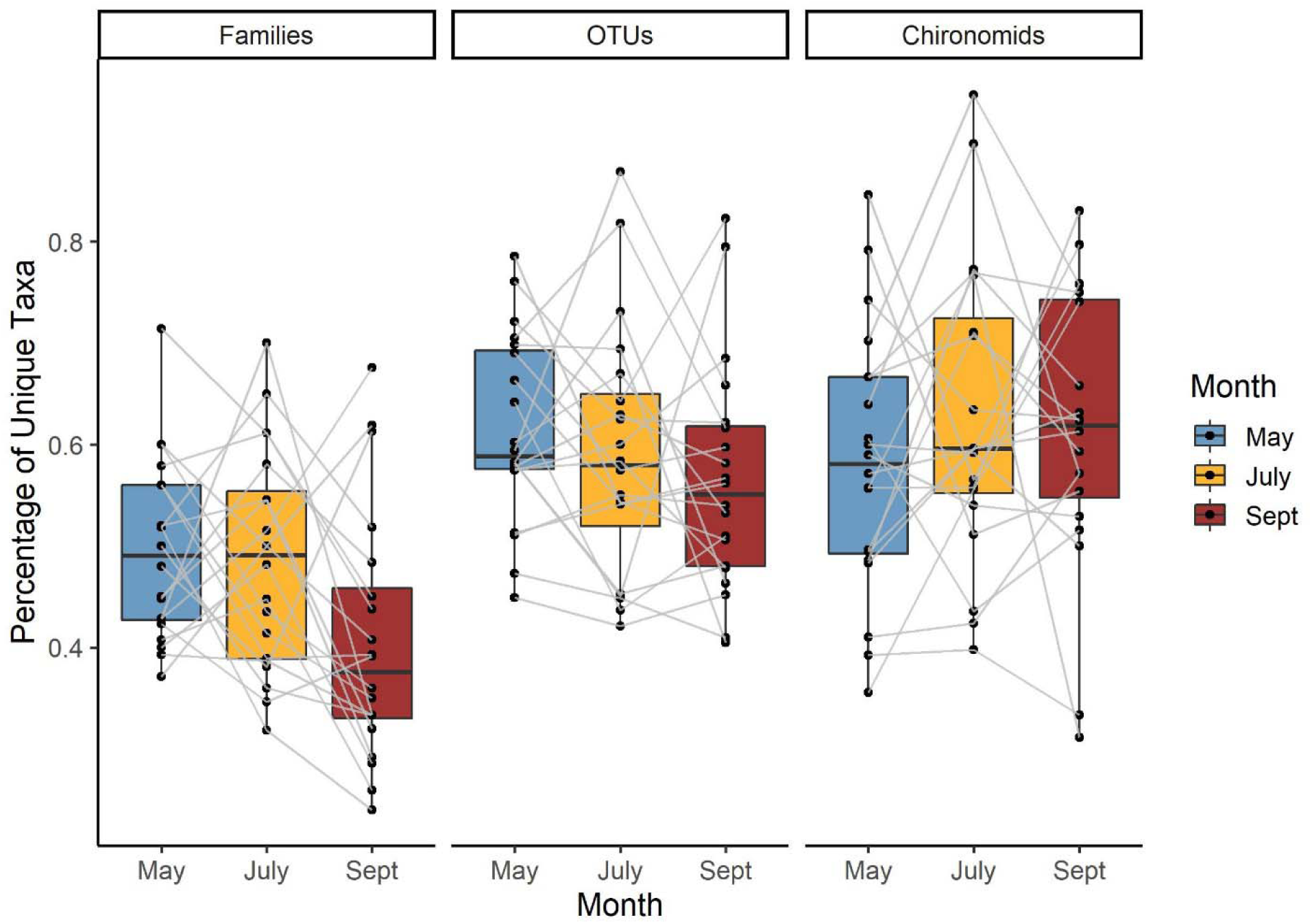
There were four biological replicates at each of the twenty streams to assess small-scale variation in community composition. Bars represent the percentage of the total number of taxa occurring at a stream that were only collected in one of four biological replicates (transects). Grey lines connect the same strean over time. There were significantly more unique or rare taxa in the OTU datasets compared to the family-level dataset (F_2,175_ = 26.1, *p* < 0.001). Sampling month had no significant effect on the percentage of rare taxa (F_2,175_ = 2.09, *p* = 0.127).

## 4. Discussion

### 4.1 Rare and missing taxa

Our approach of sampling at small spatial scales within streams revealed incredibly diverse benthic macroinvertebrate communities which varied considerably over short distances. While we collected a total of 80 samples for each sampling month, this was not sufficient to detect biodiversity at the OTU-level at either the local scale (within a stream; alpha diversity) or the regional scale (total species pool; gamma diversity). Even at family-level taxonomic resolution, we consistently underestimated total number of taxa present both locally and regionally, and this increased dramatically for OTUs. Our extrapolations suggest that after 150 samples, the accumulation curve would begin to level off for both family-level and chironomid OTUs as the estimated richness was approached; however, all invertebrate OTUs continued to increase past our extrapolation threshold. This not only indicates a vast level of diversity masked at the family level, but also suggests that we needed to double our sampling effort to represent adequately OTU-level diversity. While few aquatic metabarcoding papers explore biodiversity at the site level or compare observed versus expected number of taxa, morphological studies using family-level resolution have found similar results as our study. In shallow streams in Brazil, Ligeiro et al. (2010) detected 53 invertebrate families and determined that 81% of them were rare (in this case based on a threshold of less than 1% abundance, which differs from our definition). However, Ligeiro et al. (2010) concluded that, since sampling a single riffle in a stream collected approximately 75% of invertebrate families present, intensive sampling is not efficient and that the priority should be broad spatial coverage. In contrast, we observed that a single sample within a stream in our study is highly dissimilar in OTU composition from a second point only ten meters away and can represent anywhere between 27-85% of the total OTUs detected. Based on our results, we conclude that one sampling point is not sufficient and a more thorough sampling protocol is necessary for a more accurate representation of local diversity. While sampling intensity is an important consideration, a caveat of course is that improvements in sequencing analyses (e.g., increasing read depth) could improve the detection of rare taxa.

Our results more closely resemble terrestrial arthropod metabarcoding studies using malaise traps to explore community composition. In southern Ontario, Steinke et al. (2021a) surprisingly discovered that ten Malaise traps (tent-like insect collection traps) placed in a row had remarkably high differences in community composition. This field design of trap placement is comparable to our four kicknet transects within a stream, where we similarly found very high within-stream dissimilarity, numerous rare taxa, and large estimates of undetected taxa. For example, Steinke et al. (2021a) estimated that their OTU pool contained approximately 60% of the total diversity in the region, whereas our estimates were 69% for all invertebrate OTUs (and 78% for chironomid OTUs only). These patterns not only resemble our study in terms of missing taxa, but also display strikingly similar patterns in the number of rare taxa or those occurring only once within a site. Steinke et al. (2021a) detected close to 3000 OTUs, and almost half of these only occurred once in the same location. At the site level, the percentage of rare taxa in our study (e.g., only occurring at one of four transects in a stream) ranged from 40-90% and averaged approximately 60%.

This pattern of arthropod rarity persists across geographic ranges, as a tropical DNA barcoding study using Malaise traps (D’Souza and Hebert, 2018) observed very high beta diversity amongst traps, which did not decrease with increased local sampling, indicating that the regional species pool had not been adequately sampled. However, repeated yearly sampling of the same localities decreased beta diversity and allowed D’Souza and Hebert (2018) to determine which taxa were present yearly (even if rare) and which were ‘transient’ taxa. D’Souza and Hebert’s (2018) study clearly demonstrated the need for both spatially and temporally robust datasets to estimate accurately taxon richness and beta diversity both within a site and over time. In our dataset, we see a large proportion of rare taxa; however, repeated yearly sampling would allow us better power to determine which taxa are rare and which are transient. This is an ongoing challenge of attempting to characterize local communities in streams with no discrete boundaries and of how to delineate local sites in a continuous system. The spatial extent and sampling grain of a study can significantly alter conclusions drawn regarding community composition (Cottenie, 2005; Viana and Chase, 2019). It is possible that sampling small spatial scales like in our study can result in an overinflation of local richness due to mass effects swamping out local environmental signal (e.g., Heino et al., 2013), such as taxa being carried to the site due to water flow but not actually being able to establish there (e.g., a “sink” habitat for that species) and thus result in these transient taxa skewing dissimilarity estimates. Alternatively, sampling too large an area can miss changes in community composition in response to environmental signals due to dispersal limitation. Here, we demonstrated a vast amount of both diversity and variability in aquatic macroinvertebrates at both the local and regional scale, underscoring the importance of developing consistent, long-term monitoring programs to assess more accurately patterns in insect declines and thus better inform protection measures.

### 4.2 Taxonomic richness and land use

While both family and OTU-level richness varied between streams, this was not significantly linked to seasonality or site location (i.e., whether the stream was located on private land or on conservation authority property). This is perhaps unsurprising as local richness (or alpha diversity) can be influenced by a number of habitat parameters in aquatic systems, such as microhabitat availability in streams (Beisel et al., 1998) or hydroperiod in wetlands (Daniel et al., 2019), or can be unrelated to any habitat parameters (Ligeiro et al., 2010). While aquatic macroinvertebrates have been successfully used as indicators for decades (Buss et al., 2015; Rosenberg and Resh, 1993), it is likely that a binary category (e.g., farm or conservation area) was not an effective tool to classify stream quality, especially in a heavily impacted landscape such as southern Ontario. Streams vary in physical habitat parameters, such as the riparian buffer width and the slope of the bank, and these metrics may provide a local buffer from adjacent or upstream landscape-scale agricultural practices (Allan, 2004). The importance of local habitat conditions has been shown also in previous work in southern Ontario streams by Yates and Bailey (2010), who determined that aquatic macroinvertebrates (at family-level) were more associated with human activity directly adjacent to the stream such as channel alteration (e.g., decreased sinuosity) and buffer width. In contrast, more mobile fish communities were more responsive to landscape-scale parameters over local conditions in the same system (Yates and Bailey, 2010). Fish also have a more evident threshold response in community composition to agricultural impairment, whereas macroinvertebrates (albeit at family-level resolution) respond more gradually to such changes (Yates and Bailey, 2011) However, there is also the potential that largely developed landscapes, such as southern Ontario, have historically excluded sensitive taxa through centuries of agriculture, and invertebrate communities are already homogenous due to lack of true reference-condition streams (Krynak and Yates, 2018).

While it is possible these invertebrate communities are homogenous at the family level as suggested by Krynak and Yates (2018), and our total family count was generally in concordance with other stream studies in this region using family resolution through morphological identification (Krynak and Yates, 2018; Yates and Bailey, 2011, 2010), the total OTU richness in our study indicates that vast levels of turnover within a family are possible. At over 1600 OTUs, our metabarcoding richness appears to be much higher than stream metabarcoding studies in various geographic regions (Carew et al., 2018; Emilson et al., 2017; Kuntke et al., 2020; Serrana et al., 2019) and more closely resembles richness counts in terrestrial (Steinke et al., 2021a, 2021b) and soil (Young and Hebert, 2022) metabarcoding papers. It is likely that bioinformatic decisions in clustering and matching OTUs have a large influence on the taxonomic diversity in a dataset (Clare et al., 2016). While our choice of clustering threshold for OTU delineation (97%) is comparable to other studies, it is a more conservative choice in terms of richness than would be achieved at 98% clustering or by using exact or amplicon sequence variants (ESV/ASV; Brandt et al., 2021) and thus probably not responsible for the high level of diversity found in our dataset. Our OTU count may also be considered a conservative estimate of diversity due to the potential for taxonomic blind spots arising from the use of a single marker (COI) and primer set (Morey et al., 2020). We additionally filtered out very low-read OTUs post-clustering (less than 0.01% abundance) which would have reduced the diversity in our dataset. This filtering threshold is commonly used in other invertebrate metabarcoding studies (Beermann et al., 2018b; Steinke et al., 2021a), however it decreased the total OTUs in our dataset by half. However, while these OTUs made up 50% of the total OTUs, they only corresponded to 33,840 sequence reads (0.06% of total sequences) and it is important to define a threshold to eliminate OTUs based upon erroneous sequences. We also selected a threshold of 90% sequence similarity to the reference database (BOLD) in order to be included in the final dataset. While many studies use a conservative 98% threshold (e.g., Emilson et al., 2017; Steinke et al., 2021), others have selected a 85% similarity threshold (e.g., Kuntke et al., 2020). In our dataset, it is likely that 98% is too strict a cut-off due to the sparsity of reference sequences for understudied taxa such as chironomids. A match of 90% similarity is generally accepted as a ‘family-level’ match in multiple metabarcoding studies (e.g., Emilson et al., 2017; Kuntke et al., 2020) and thus aligned well with our approach of comparing OTU and family-level diversity.

In our system, chironomids had the highest OTU count of any family. This is consistent with previous aquatic metabarcoding work, notably by Beermann et al. (2018b), who detected nearly 200 chironomid OTUs in a stream mesocosm experiment. Of these total chironomid OTUs, 85% did not have a binomial name assigned from BOLD, which is nearly identical to the value we detected (84.8% without a binomial name in the reference database, high similarity matchs). Even with unnamed OTUs, Beerman et al. (2018b) detected unique responses to environmental stressors (even amongst OTUs which matched to the same binomial name). Reference databases are not complete for many groups, especially very species-rich groups or difficult-to-identify taxa such as aquatic insect larvae, and improvement in species coverages must be made in order to improve the quality of metabarcoding datasets. It is also important to consider that the use of 97% as a clustering threshold can also result in the “splitting” of OTUs (e.g., a single species with large intraspecific divergence of CO1 being split into two OTUs) and thus artificially inflating the OTU count. For example, some chironomid species complexes can have as high as a ten percent divergence (Lin et al., 2017), which would result in multiple OTU clusters when using a 97% threshold. Ultimately, there are numerous bioinformatic decisions which can either increase or decrease the number of OTUs in a metabarcoding study. The vast diversity of the chironomid family in particular merits future study to determine the extent of haplotype diversity which may interfere with establishing discrete OTUs using a set clustering threshold (e.g., Beermann et al., 2018b; Lin et al., 2017).

### 4.3 Dissimilarity and habitat heterogeneity

Mean Raup-Crick dissimilarity values were consistently high for all levels of taxonomic assignment and sampling season, indicating that our biological replicates were quite unique from each other even when collected at the same stream. While a categorical land use variable did not prove informative in distinguishing macroinvertebrate communities from farm or conservation area streams, we expected the percentage of land used for cropping in the catchment would be a more accurate indication of stream condition. We expected an increased level of agriculture would result in more homogeneous invertebrate communities within a stream (i.e., lower beta diversity) due to the reduction in habitat complexity (e.g., as in Astorga et al., 2014; Maloney et al., 2011; Martins et al., 2017; Siqueira et al., 2015). We detected no significant relationship between land use and beta diversity (as in Heino et al., 2013; Petsch et al., 2021), and dissimilarity values were quite variable.

Interestingly, we observed lower dissimilarity values in May for all groups compared to other sampling months. While it is possible this is a spurious correlation, Zizka et al. (2020) observed seasonal community changes for aquatic macroinvertebrate OTUs in a study of urban streams. In reference condition (“near natural”) streams, Zizka et al. (2020) discovered that the community composition of invertebrates stayed consistent across seasonal sampling periods, whereas communities in more highly impacted streams differed across seasonal sampling periods. This contrasting pattern in seasonal change may be in response to the fact that the inflow of stressors can change over time in impacted areas, resulting in a community-level response that differs over time (Zizka et al., 2020). It is possible here that changes in water quality or other habitat parameters during the spring resulted in more homogenous communities compared to later in the sampling season. Of course, this may also be due to the emergence of different insect taxa in streams over a season. Our results here indicate that the percentage of agriculture in the landscape is not sufficient here to detect a complicated influence of land use; more detailed parameters are necessary, or our sampling was not sufficient to detect any homogenization, or perhaps the variation in land use in our study was not significant enough to detect a gradient response in community composition.

### 4.4 mplications

Metabarcoding revealed a huge amount of diversity in southern Ontario streams, with many rare or undetected taxa present within each stream. This study highlights the importance of developing long-term monitoring plans, especially for ecological indicator taxa such as aquatic macroinvertebrate taxa. Field sampling design is an extremely important consideration in monitoring studies, and by using molecular identification we reveal that one sampling point in a stream in not sufficient to detect local diversity. While we detected no discernable response in taxonomic richness or within-stream dissimilarity to agricultural land use, it is possible our interpretation of agriculture land use was too narrow. Given the importance of local parameters to aquatic invertebrate communities, future metabarcoding studies should consider more precise estimates of land use, including both water chemistry and physical parameters, such as stream sinuosity, riparian buffer width, and bank slope to reflect more accurately site condition. In general, our conclusions regarding OTU diversity and rarity were similar to terrestrial metabarcoding work and indicate that aquatic biomonitoring programs can benefit from molecular identification to more accurately reflect trends in insect biodiversity.

## Supporting information

Appendix 1 - Data cleaning methods

## Data availability

Sequencing data will be made available at NCBI Sequence Read Archive upon publication.

## Author contributions

JEG, RH, and KC conceived the ideas; JEG designed the study, performed the field and laboratory work, performed the bioinformatics and data analysis, and wrote the first draft of the manuscript; JEG, RH, and KC edited the manuscript.

## Acknowledgements

We would like to acknowledge that our stream sampling sites are located on the traditional territory of the Mississaugas of the Credit First Nations, the Anishnabek, Haudenosaunee (Iroquis), and Ojibway/Chippewa peoples.

We would like to thank Marie Gutgesell, Emily Champagne, Christine Dulal-Whiteway, and Dr. Kevin McCann for their assistance locating study sites and Evan Versteeg for assistance calculating land use in ArcGIS. We would also like to thank Ian Thompson for his valuable assistance collecting and sorting benthic macroinvertebrate samples, Dr. Yoamel Milián-García for helpful advice and feedback throughout lab work, and Dr. Sarah (Sally) Adamowicz for helpful suggestions regarding our bioinformatics pipeline. Thank you to staff at the following conservation authorities: Catfish Creek CA, Kettle Creek CA, Long Point CA, Nature Conservancy Canada and Thames Talbot Land Trust.

## Supplementary Information

**Raw sequencing data:** Available on NCBI’s SRA under PRJNA783201.

**Appendix 1 –** A word document elaborating on data cleaning protocols post-bioinformatic processing using metabaR.

**Appendix 2** – The R code used to clean the data following the protocol in Appendix 1.

**Appendix 3** – Raw data post JAMP processing, to be used in Appendix 3. Composed of 4 csv files (Read table, OTU taxonomy data, PCR metadata, sample metadata).

**Appendix 4** – The R code used to analysis the data and generate figures. This code is meant to run using the output from Appendix 2.

## References

Albert, J.S., Destouni, G., Duke-Sylvester, S.M., Magurran, A.E., Oberdorff, T., Reis, R.E., Winemiller, K.O., Ripple, W.J., 2021. Scientists’ warning to humanity on the freshwater biodiversity crisis. Ambio 50, 85–94. https://doi.org/10.1007/s13280-020-01318-8

Allan, J.D., 2004. Landscapes and riverscapes: The influence of land use on stream ecosystems. Annu. Rev. Ecol. Evol. Syst. 35, 257–284. https://doi.org/10.1146/annurev.ecolsys.35.120202.110122

Astorga, A., Death, R., Death, F., Paavola, R., Chakraborty, M., Muotka, T., 2014. Habitat heterogeneity drives the geographical distribution of beta diversity: The case of New Zealand stream invertebrates. Ecol. Evol. 4, 2693–2702. https://doi.org/10.1002/ece3.1124

Baird, D.J., Hajibabaei, M., 2012. Biomonitoring 2.0: A new paradigm in ecosystem assessment made possible by next-generation DNA sequencing. Mol. Ecol. 21, 2039–2044. https://doi.org/10.1111/j.1365-294X.2012.05519.x

Beermann, A.J., Elbrecht, V., Karnatz, S., Ma, L., Matthaei, C.D., Piggott, J.J., Leese, F., 2018a. Multiple-stressor effects on stream macroinvertebrate communities: A mesocosm experiment manipulating salinity, fine sediment and flow velocity. Sci. Total Environ. 610– 611, 961–971. https://doi.org/10.1016/j.scitotenv.2017.08.084

Beermann, A.J., Zizka, V.M.A., Elbrecht, V., Baranov, V., Leese, F., 2018b. DNA metabarcoding reveals the complex and hidden responses of chironomids to multiple stressors. Environ. Sci. Eur. 30, 26. https://doi.org/10.1186/s12302-018-0157-x

Beisel, J.N., Usseglio-Polatera, P., Thomas, S., Moreteau, J.C., 1998. Stream community structure in relation to spatial variation: The influence of mesohabitat characteristics. Hydrobiologia 389, 73–88. https://doi.org/10.1023/A:1003519429979

Blackman, R., Mächler, E., Altermatt, F., Arnold, A., Beja, P., Boets, P., Egeter, B., Elbrecht, V., Filipe, A.F., Jones, J., Macher, J., Majaneva, M., Martins, F., Múrria, C., Meissner, K., Pawlowski, J., Schmidt Yáñez, P., Zizka, V., Leese, F., Price, B., Deiner, K., 2019. Advancing the use of molecular methods for routine freshwater macroinvertebrate biomonitoring – the need for calibration experiments. Metabarcoding and Metagenomics 3. https://doi.org/10.3897/mbmg.3.34735

Brandt, M.I., Günther, B., Arnaud-Haond, S., 2021. Bioinformatic pipelines combining denoising and clustering tools allow for more comprehensive prokaryotic and eukaryotic metabarcoding. Mol. Ecol. Resour. https://doi.org/10.1016/s0740-5472(96)90021-5

Brooks, R.T., 2000. Annual and Seasonal Variation and the Effects of Hydroperiod on Benthic Macroinvertebrates of Seasonal Forest (“Vernal”) Ponds in Central Massachusetts, USA. Wetlands 20, 707–715. http://dx.doi.org/10.1672/0277-5212(2000)020[0707:AASVAT]2.0.CO;2

Buchner, D., Leese, F., 2020. BOLDigger -a Python package to identify and organise sequences with the Barcode of Life Data systems. Metabarcoding and Metagenomics 4, 19–21. https://doi.org/10.3897/mbmg.4.53535

Buss, D.F., Carlisle, D.M., Chon, T.-S., Culp, J., Harding, J.S., Keizer-Vlek, H.E., Robinson, W.A., Strachan, S., Thirion, C., Hughes, R.M., 2015. Stream biomonitoring using macroinvertebrates around the globe: a comparison of large-scale programs. Environ. Monit. Assess. 187. https://doi.org/10.1007/s10661-014-4132-8

Carew, M.E., Kellar, C.R., Pettigrove, V.J., Hoffmann, A.A., 2018. Can high-throughput sequencing detect macroinvertebrate diversity for routine monitoring of an urban river? Ecol. Indic. 85, 440–450. https://doi.org/10.1016/j.ecolind.2017.11.002

Ceballos, G., Ehrlich, P.R., Barnosky, A.D., García, A., Pringle, R.M., Palmer, T.M., 2015. Accelerated modern human-induced species losses: Entering the sixth mass extinction. Sci. Adv. 1, 9–13. https://doi.org/10.1126/sciadv.1400253

Ceballos, G., Ehrlich, P.R., Dirzo, R., 2017. Biological annihilation via the ongoing sixth mass extinction signaled by vertebrate population losses and declines. Proc. Natl. Acad. Sci. U. S. A. 114, E6089–E6096. https://doi.org/10.1073/pnas.1704949114

Ceballos, G., Ehrlich, P.R., Raven, P.H., 2020. Vertebrates on the brink as indicators of biological annihilation and the sixth mass extinction. Proc. Natl. Acad. Sci. U. S. A. 117, 13596–13602. https://doi.org/10.1073/pnas.1922686117

Clare, E.L., Chain, F.J.J., Littlefair, J.E., Cristescu, M.E., Deiner, K., 2016. The effects of parameter choice on defining molecular operational taxonomic units and resulting ecological analyses of metabarcoding data. Genome 59, 981–990. https://doi.org/10.1139/gen-2015-0184

Costa, S.S., Melo, A.S., 2008. Beta diversity in stream macroinvertebrate assemblages: Among-site and among-microhabitat components. Hydrobiologia 598, 131–138. https://doi.org/10.1007/s10750-007-9145-7

Costello, M.J., May, R.M., Stork, N.E., 2013. Can we name earth’s species before they go extinct? Science 339, 413–416. https://doi.org/10.1126/science.1230318

Cottenie, K., 2005. Integrating environmental and spatial processes in ecological community dynamics. Ecol. Lett. 8, 1175–1182. https://doi.org/10.1111/j.1461-0248.2005.00820.x

Cristescu, M.E., 2014. From barcoding single individuals to metabarcoding biological communities: Towards an integrative approach to the study of global biodiversity. Trends Ecol. Evol. 29, 566–571. https://doi.org/10.1016/j.tree.2014.08.001

D’Souza, M.L., Hebert, P.D.N., 2018. Stable baselines of temporal turnover underlie high beta diversity in tropical arthropod communities. Mol. Ecol. 27, 2447–2460. https://doi.org/10.1111/mec.14693

Daniel, J., Gleason, J.E., Cottenie, K., Rooney, R.C., 2019. Stochastic and deterministic processes drive wetland community assembly across a gradient of environmental filtering. Oikos 1158–1169. https://doi.org/10.1111/oik.05987

Dias-Silva, K., Brasil, L.S., Veloso, G.K.O., Cabette, H.S.R., Juen, L., 2020. Land use change causes environmental homogeneity and low beta-diversity in Heteroptera of streams. Ann. Limnol. 56. https://doi.org/10.1051/limn/2020007

Dobson, M., 1994. Microhabitat as a determinant of diversity: stream invertebrates colonizing leaf packs. Freshw. Biol. 32, 565–572. https://doi.org/10.1111/j.1365-2427.1994.tb01147.x

Dudgeon, D., 2019. Multiple threats imperil freshwater biodiversity in the Anthropocene. Curr. Biol. 29, R960–R967. https://doi.org/10.1016/j.cub.2019.08.002

Edgar, R.C., 2010. Search and clustering orders of magnitude faster than BLAST. Bioinformatics. 26, 2460–2461. https://doi.org/10.1093/bioinformatics/btq461

Elbrecht, V., Leese, F., 2017. Validation and development of COI metabarcoding primers for freshwater macroinvertebrate bioassessment. Front. Environ. Sci. 5, 1–11. https://doi.org/10.3389/fenvs.2017.00011

Elbrecht, V., Vamos, E.E., Meissner, K., Aroviita, J., Leese, F., 2017a. Assessing strengths and weaknesses of DNA metabarcoding-based macroinvertebrate identification for routine stream monitoring. Methods Ecol. Evol. 1265–1275. https://doi.org/10.1111/2041-210X.12789

Elbrecht, V., Vamos, E.E., Meissner, K., Aroviita, J., Leese, F., 2017b. Assessing strengths and weaknesses of DNA metabarcoding-based macroinvertebrate identification for routine stream monitoring. Methods Ecol. Evol. 8, 1265–1275. https://doi.org/10.1111/2041-210X.12789

Emilson, C.E., Thompson, D.G., Venier, L.A., Porter, T.M., Swystun, T., Chartrand, D., Capell, S., Hajibabaei, M., 2017. DNA metabarcoding and morphological macroinvertebrate metrics reveal the same changes in boreal watersheds across an environmental gradient. Sci. Rep. 7, 1–12. https://doi.org/10.1038/s41598-017-13157-x

Esri, 2020. ArcGIS. Esri Inc. https://www.esri.com/en-us/arcgis/products/arcgis-pro/overview.

Fourcade, Y., Åström, S., Öckinger, E., 2019. Climate and land-cover change alter bumblebee species richness and community composition in subalpine areas. Biodivers. Conserv. 28, 639–653. https://doi.org/10.1007/s10531-018-1680-1

Gleason, J., Rooney, R., 2017. Aquatic invertebrates are poor indicators of land use in northern prairie pothole region wetlands. Ecol. Indic. 81, 333–337. https://doi.org/10.1016/j.ecolind.2017.06.013

Gleason, J.E., Elbrecht, V., Braukmann, T.W.A., Hanner, R.H., Cottenie, K., 2020. Assessment of stream macroinvertebrate communities with eDNA is not congruent with tissue-based metabarcoding. Mol. Ecol. 1–13. https://doi.org/10.1111/mec.15597

Government of Ontario, 2020. User Guide for Ontario Flow Assessment Tool. Ministry of Northern Development, Mines, Natural Resources and Forestry. 53 pages.

Grant, E.H.C., Lowe, W.H., Fagan, W.F., 2007. Living in the branches: Population dynamics and ecological processes in dendritic networks. Ecol. Lett. 10, 165–175. https://doi.org/10.1111/j.1461-0248.2006.01007.x

Haase, P., Murray-Bligh, J., Lohse, S., Pauls, S., Sundermann, A., Gunn, R., Clarke, R., 2006. Assessing the impact of errors in sorting and identifying macroinvertebrate samples. Hydrobiologia 566, 505–521. https://doi.org/10.1007/s10750-006-0075-6

Haase, P., Pauls, S.U., Schindehu, K., Sundermann, A., 2010. First audit of macroinvertebrate samples from an EU Water Framework Directive monitoring programL: human error greatly lowers precision of assessment results. J. North Am. Benthol. Soc. 29, 1279–1291. https://doi.org/10.1899/09-183.1

Hallmann, C.A., Sorg, M., Jongejans, E., Siepel, H., Hofland, N., Schwan, H., Stenmans, W., Müller, A., Sumser, H., Hörren, T., Goulson, D., De Kroon, H., 2017. More than 75 percent decline over 27 years in total flying insect biomass in protected areas. PLoS One 12. https://doi.org/10.1371/journal.pone.0185809

Heino, J., Grönroos, M., Ilmonen, J., Karhu, T., Niva, M., Paasivirta, L., 2013. Environmental heterogeneity and β diversity of stream macroinvertebrate communities at intermediate spatial scales. Freshw. Sci. 32, 142–154. https://doi.org/10.1899/12-083.1

Heino, J., Louhi, P., Muotka, T., 2004. Identifying the scales of variability in stream macroinvertebrate abundance, functional composition and assemblage structure. Freshw. Biol. 49, 1230–1239. https://doi.org/10.1111/j.1365-2427.2004.01259.x

Hsieh, T., Ma, K., Chao, A., 2020. iNEXT: Interpolation and Extrapolation for Species Diversity. R package version 2.0.20, http://chao.stat.nthu.edu.tw/wordpress/software_download/.

Jones, F.C., Somers, K.M., Craig, B., Reynoldson, T.B., 2007. Ontario Benthos Biomonitoring Network: Protocol Manual. Ontario Ministry of Environment, Dorset, ON, 109 pages.

Krynak, E.M., Yates, A.G., 2018. Benthic invertebrate taxonomic and trait associations with land use in an intensively managed watershed: Implications for indicator identification. Ecol. Indic. 93, 1050–1059. https://doi.org/10.1016/j.ecolind.2018.06.002

Kuntke, F., de Jonge, N., Hesselsøe, M., Lund Nielsen, J., 2020. Stream water quality assessment by metabarcoding of invertebrates. Ecol. Indic. 111, 105982. https://doi.org/10.1016/j.ecolind.2019.105982

Ligeiro, R., Melo, A.S., Callisto, M., 2010. Spatial scale and the diversity of macroinvertebrates in a Neotropical catchment. Freshw. Biol. 55, 424–435. https://doi.org/10.1111/j.1365-2427.2009.02291.x

Lin, X., Stur, E., Ekrem, T., 2018. DNA barcode and morphology reveal unrecognized species in Chironomidae (Diptera). Insect Sys. & Evo. 49, 329–398. https://doi.org/10.1163/1876312X-00002172

Maggia, M.E., Decaëns, T., Lapied, E., Dupont, L., Roy, V., Schimann, H., Orivel, J., Murienne, J., Baraloto, C., Cottenie, K., Steinke, D., 2021. At each site its diversity: DNA barcoding reveals remarkable earthworm diversity in neotropical rainforests of French Guiana. Appl. Soil Ecol. 164. https://doi.org/10.1016/j.apsoil.2021.103932

Maloney, K.O., Munguia, P., Mitchell, R.M., 2011. Anthropogenic disturbance and landscape patterns affect diversity patterns of aquatic benthic macroinvertebrates. J. North Am. Benthol. Soc. 30, 284–295. https://doi.org/10.1899/09-112.1

Martin, M., 2011. Cutadapt removes adapter sequences from high-throughput sequencing reads. Embnet Journal. 17, 10–12. https://doi.org/10.14806/ej.17.1.200

Martins, R.T., Couceiro, S.R.M., Melo, A.S., Moreira, M.P., Hamada, N., 2017. Effects of urbanization on stream benthic invertebrate communities in Central Amazon. Ecol. Indic. 73, 480–491. https://doi.org/10.1016/j.ecolind.2016.10.013

Milián-García, Y., Young, R., Madden, M., Bullas-Appleton, E., Hanner, R.H., 2021. Optimization and validation of a cost-effective protocol for biosurveillance of invasive alien species. Ecol. Evol. 11, 1999–2014. https://doi.org/10.1002/ece3.7139

Montgomery, G.A., Dunn, R.R., Fox, R., Jongejans, E., Leather, S.R., Saunders, M.E., Shortall, C.R., Tingley, M.W., Wagner, D.L., 2020. Is the insect apocalypse upon us? How to find out. Biol. Conserv. 241. https://doi.org/10.1016/j.biocon.2019.108327

Mora, C., Tittensor, D.P., Adl, S., Simpson, A.G.B., Worm, B., 2011. How many species are there on earth and in the ocean? PLoS Biol. 9, 1–8. https://doi.org/10.1371/journal.pbio.1001127

Morey, K.C., Bartley, T.J., Hanner, R.H., 2020. Validating environmental DNA metabarcoding for marine fishes in diverse ecosystem using a public aquarium. Environmental DNA. 2, 330–342. https://doi.org/10.1002/edn3.76

Nicacio, G., Juen, L., 2015. Chironomids as indicators in freshwater ecosystems: An assessment of the literature. Insect Conserv. Divers. 8, 393–403. https://doi.org/10.1111/icad.12123

Oksanen, J., Blanchet, F.G., Kindt, R., Legendre, P., O’Hara, R.B., Simpson, G.L., Solymos, P., Stevens, M.H.H., Wagner, H., 2017. vegan: Community Ecology Package. R Packag. version 2.4-2. https://cran.r-project.org/web/packages/vegan

Ontario Ministry of Natural Resources and Forestry, 2016. Ontario Land Cover Compilation Data Specifications Version 2 .0. https://www.sse.gov.on.ca/sites/MNR-PublicDocs/EN/CMID/Ontario Land Cover Compilation -Data Specification Version.pdf

Ortega, J.C.G., Thomaz, S.M., Bini, L.M., 2018. Experiments reveal that environmental heterogeneity increases species richness, but they are rarely designed to detect the underlying mechanisms. Oecologia 188, 11–22. https://doi.org/10.1007/s00442-018-4150-2

Pauls, S.U., Alp, M., Bálint, M., Bernabò, P., Čiampor, F., Čiamporová-Zaťovičová, Z., Finn, D.S., Kohout, J., Leese, F., Lencioni, V., Paz-Vinas, I., Monaghan, M.T., 2014. Integrating molecular tools into freshwater ecology: Developments and opportunities. Freshw. Biol. 59, 1559–1576. https://doi.org/10.1111/fwb.12381

Persaud, S.F., Cottenie, K., Gleason, J.E., 2021. Ethanol eDNA reveals unique community composition of aquatic macroinvertebrates compared to bulk tissue metabarcoding in a biomonitoring sampling scheme. Diversity 13, 1–15. https://doi.org/10.3390/d13010034

Petsch, D.K., Saito, V.S., Landeiro, V.L., Silva, T.S.F., Bini, L.M., Heino, J., Soininen, J., Tolonen, K.T., Jyrkänkallio-Mikkola, J., Pajunen, V., Siqueira, T., Melo, A.S., 2021. Beta diversity of stream insects differs between boreal and subtropical regions, but land use does not generally cause biotic homogenization. Freshw. Sci. 40, 53–64. https://doi.org/10.1086/712565

QGISDevelopmentTeam, 2020. QGIS Geographic Information System. Open Source Geospatial Foundation Project. http://qgis.osgeo.org“.

R Core Team, 2020. R: A language and environment for statistical computing, version 4.03.Vienna, Austria: R Foundation for Statistical computing.

Ratnasingham, S., Hebert, P.D.N., 2007. BOLD: The Barcode of Life Data System (www.barcodinglife.org). Mol. Ecol. Notes 7, 355–364. https://doi.org/10.1111/j.1471-8286.2006.01678.x

Raven, P.H., Wagner, D.L., 2021. Agricultural intensification and climate change are rapidly decreasing insect biodiversity. Proc. Natl. Acad. Sci. U. S. A. 118, 1–6. https://doi.org/10.1073/PNAS.2002548117

Reid, A.J., Carlson, A.K., Creed, I.F., Eliason, E.J., Gell, P.A., Johnson, P.T.J., Kidd, K.A., MacCormack, T.J., Olden, J.D., Ormerod, S.J., Smol, J.P., Taylor, W.W., Tockner, K., Vermaire, J.C., Dudgeon, D., Cooke, S.J., 2019. Emerging threats and persistent conservation challenges for freshwater biodiversity. Biol. Rev. 94, 849–873. https://doi.org/10.1111/brv.12480

Rosenberg, D.M., Resh, V.H., 1993. Freshwater biomonitoring and benthic invertebrates. Chapman & Hall, New York, NY.

Rosenberg, K. V., Dokter, A.M., Blancher, P.J., Sauer, J.R., Smith, A.C., Smith, P.A., Stanton, J.C., Panjabi, A., Helft, L., Parr, M., Marra, P.P., 2019. Decline of the North American avifauna. Science. 366, 120–124. https://doi.org/10.1126/science.aaw1313

Sánchez-Bayo, F., Wyckhuys, K.A.G., 2019. Worldwide decline of the entomofauna: A review of its drivers. Biol. Conserv. 232, 8–27. https://doi.org/10.1016/j.biocon.2019.01.020

Serrana, J.M., Miyake, Y., Gamboa, M., Watanabe, K., 2019. Comparison of DNA metabarcoding and morphological identification for stream macroinvertebrate biodiversity assessment and monitoring. Ecol. Indic. 101, 963–972. https://doi.org/10.1016/j.ecolind.2019.02.008

Siqueira, T., Lacerda, C.G.L.T., Saito, V.S., 2015. How Does Landscape Modification Induce Biological Homogenization in Tropical Stream Metacommunities? Biotropica 47, 509–516. https://doi.org/10.1111/btp.12224

Stanfield, L. (editor), 2017. Ontario Stream Assessment Protocol. Version 8.0. Fisheries Policy Section. Ontario Ministry of Natural Resources. Peterborough, Ontario. 376 pages.

Stein, A., Gerstner, K., Kreft, H., 2014. Environmental heterogeneity as a universal driver of species richness across taxa, biomes and spatial scales. Ecol. Lett. 17, 866–880. https://doi.org/10.1111/ele.12277

Steinke, D., Braukmann, T.W., Manerus, L., Woodhouse, A., Elbrecht, V., 2021a. Effects of Malaise trap spacing on species richness and composition of terrestrial arthropod bulk samples. Metabarcoding and Metagenomics 5, 43–50. https://doi.org/10.3897/mbmg.5.59201

Steinke, D., DeWaard, S., Sones, J., Ivanova, N. V., Prosser, S.W.J., Perez, K., Braukmann, T.W.A., Milton, M., Zakharov, E. V., deWaard, J.R., Ratnasingham, S., Hebert, P.D.N., 2021b. Message in a Bottle – Metabarcoding Enables Biodiversity Comparisons Across Ecoregions. bioRxiv.

Taberlet, P., Coissac, E., Pompanon, F., Brochmann, C., Willerslev, E., 2012. Towards next-generation biodiversity assessment using DNA metabarcoding. Mol. Ecol. 21, 2045–2050. https://doi.org/10.1111/j.1365-294X.2012.05470.x

Thomas, C.D., Jones, T.H., Hartley, S.E., 2019. “Insectageddon”: A call for more robust data and rigorous analyses. Glob. Chang. Biol. 25, 1891–1892. https://doi.org/10.1111/gcb.14608

Thomas, J.A., 2005. Monitoring change in the abundance and distribution of insects using butterflies and other indicator groups. Philos. Trans. R. Soc. B Biol. Sci. 360, 339–357. https://doi.org/10.1098/rstb.2004.1585

Viana, D.S., Chase, J.M., 2019. Spatial scale modulates the inference of metacommunity assembly processes. Ecology 100, e02576. https://doi.org/10.1002/ecy.2576

Wagner, D.L., Grames, E.M., Forister, M.L., Berenbaum, M.R., Stopak, D., 2021. Insect decline in the Anthropocene: Death by a thousand cuts. Proc. Natl. Acad. Sci. U. S. A. 118, 1–10. https://doi.org/10.1073/PNAS.2023989118

Wickham, H., 2020. ggplot2: Elegant Graphics for Data Analysis. Springer-Verlag New York. ISBN 978-3-319-24277-4, https://ggplot2.tidyverse.org

Yates, A.G., Bailey, R.C., 2011. Effects of taxonomic group, spatial scale and descriptor on the relationship between human activity and stream biota. Ecol. Indic. 11, 759–771. https://doi.org/10.1016/j.ecolind.2010.09.003

Yates, A.G., Bailey, R.C., 2010. Covarying patterns of macroinvertebrate and fish assemblages along natural and human activity gradients: Implications for bioassessment. Hydrobiologia 637, 87–100. https://doi.org/10.1007/s10750-009-9987-2

Young, M.R., Hebert, P.D.N., 2022. Unearthing soil arthropod diversity through DNA metabarcoding. PeerJ. 10:e12845. https://doi.org/10.7717/peerj.12845

Zinger, L., Bonin, A., Alsos, I.G., Bálint, M., Bik, H., Boyer, F., Chariton, A.A., Creer, S., Coissac, E., Deagle, B.E., De Barba, M., Dickie, I.A., Dumbrell, A.J., Ficetola, G.F., Fierer, N., Fumagalli, L., Gilbert, M.T.P., Jarman, S., Jumpponen, A., Kauserud, H., Orlando, L., Pansu, J., Pawlowski, J., Tedersoo, L., Thomsen, P.F., Willerslev, E., Taberlet, P., 2019. DNA metabarcoding—Need for robust experimental designs to draw sound ecological conclusions. Mol. Ecol. mec.15060. https://doi.org/10.1111/mec.15060

Zinger, L., Lionnet, C., Benoiston, A.S., Donald, J., Mercier, C., Boyer, F., 2021. metabaR: An r package for the evaluation and improvement of DNA metabarcoding data quality. Methods Ecol. Evol. 2021, 1–7. https://doi.org/10.1111/2041-210X.13552

Zizka, V.M.A., Geiger, M.F., Leese, F., 2020. DNA metabarcoding of stream invertebrates reveals spatio-temporal variation but consistent status class assessments in a natural and urban river. Ecol. Indic. 115, 106383. https://doi.org/10.1016/j.ecolind.2020.106383

