## Appendix 1 - Data cleaning methods for "Hidden diversity – DNA metabarcoding reveals hyper-diverse benthic invertebrate communities"

**Appendix 1 – Data quality control methods**

**Title: Hidden diversity – DNA metabarcoding reveals hyper-diverse benthic invertebrate communities**

**Author names and affiliations:** Jennifer Erin Gleason^1,*^, Robert Hanner^1^, Karl Cottenie^1^

^1^ Department of Integrative Biology, University of Guelph, Guelph, Ontario, Canada

^*^ corresponding author

We used the R package metabaR (v. 1.0.0; Zinger et al., 2021) to assess the quality of our metabarcoding data. The code used to analyze data quality was adapted from the metabaR tutorial available on github: (https://github.com/metabaRfactory/metabaR/blob/master/vignettes/metabaRF-vignette.Rmd) and our adapted version for this dataset is available as supplementary information. All functions described below are from the metabaR package unless otherwise stated. Plots were created using ggplot2 (v. 3.3.3; Wickham, 2009).

We first plotted the total number of reads and the total number of OTUs in each negative control type (extraction, PCR, sequencing) and samples to visualize the proportion of reads/OTUs in each control type (Figure 1). To visually assess that sequencing depth was sufficient, we plotted the number of sequence reads per PCR reaction and the number of OTUs to confirm that there was no correlation between read number and OTU count (Figure 2). If reads were significantly correlated with OTU number, it would indicate that sequencing depth was not sufficient. We used the function ‘ggpcrplate’ to visualize the number of reads per PCR reaction between samples and controls in plate format to confirm that there was no pattern in samples containing low sequence reads (e.g., all in the same row/column; Figure 3). To assess sequencing depth in each sample, we used the function ‘hill_rarefaction’ with a bootstrap value (the number of resampling steps) of 20. Rarefaction curves were generated using species richness, Shannon index, inverted Simpon’s index (Hill numbers of q = 0, 1, 2 respectively; Chao et al., 2014) and Good’s coverage index (Figure 4).

To flag potential contaminants, we first subset the data into the four respective plates using the function ‘subset_metabarlist’. As each plate was processed and sequenced separately, the negative controls were unique to a plate. For each plate, we used the function ‘constaslayer’ to identify and flag OTUs as contaminants if the relative abundance of that OTU across the entire dataset was highest in a negative control. For a sample to be excluded as contaminated, more than 10% of reads within that sample had to correspond to an OTU flagged as a contaminate. No samples were removed due to contamination as this threshold as not met.

We flagged OTUs as non-target amplification if they did not belong to the phyla we expected to be present in our dataset: arthropods, molluscs and annelids. We chose not to include nematodes here as our dataset contained very few nematode sequence reads (4397). This could be due to their potentially small size they were not sufficiently collected in the field or recognized during sample sorting. The majority of sequences corresponding to nematodes matched to *Gordius* sp. (3929), which are large horsehair worms that would not have been missed during collection/sorting. It is also possible that nematode sequences were not amplified effectively due to primer biases. There were 40,620,884 arthropod sequences, 9,055,936 annelida sequences and 1,166,259 mollusc sequences. In addition to flagged OTUs that were not our target taxa, we also flagged those that had a match similarity of less than 90% to our reference database (Figure 5). While some studies use a conservative 98% threshold (e.g., Emilson et al., 2017; Steinke et al., 2021), others have selected a 85% similarity threshold (e.g., Kuntke et al., 2020). In our dataset, it is likely that 98% is too strict a cut-off due to the sparsity of reference sequences for understudied taxa such as chironomids. We based our threshold on examining the spread of our similarity data (Figure 5), and selected 90% to balance the inclusion of species not recorded on the Barcode of Life Database (BOLD; Ratnasingham and Hebert, 2007) while still excluding potentially low quality sequences. The average sequence similarity for our target taxa was 96%.

Based on our initial three sequencing runs, we calculated the average and standard deviation of sequence reads for all samples (153,632 ± 66,288). We re-ran any samples that contained fewer reads than one standard deviation below the average (less than 87,344 sequences), beginning from the extraction step to ensure no errors were carried over from previous laboratory work. This ‘redo plate’ was then sequenced and all OTUs were clustered together to create the final dataset. When filtering data, we retained this cut-off threshold for sequence reads and excluded samples that had too little sequence reads (Figure 6). In total, this only removed three biological replicates out of 240 in our dataset.

To assess how consistent PCR replicates were, we calculated pairwise Bray-Curtis dissimilarity within samples (e.g., between technical replicates) and between samples using the function ‘pcr_within_between’ and visualized the intersection of these values using ‘check_pcr_thresh’ (Figure 7). We expect PCR replicates that perform well will be more similar to each other than to other samples (i.e., a low Bray-Curtis dissimilarity value). We used the threshold of intersection (0.51) as our cut-off value to remove any outlier PCRs with within-sample dissimilarity scores higher than this. This means a sample would be flagged if the distance within a sample (e.g., between technical replicates) was greater than expected given the typical dissimilarity within and between samples.

Tag-jumps occur when a sequence is erroneously assigned to a sample it does not belong to, resulting in a false positive. The frequency of tag-jumps in a dataset can be assessed using sequencing negative controls. We used the function ‘tagjumpslayer’, which reduces an OTU abundance in a sample to zero if the total abundance (i.e., sequence reads) of that OTU across the entire dataframe is lower than a set threshold. This threshold is when the relative abundance of an OTU in a sample is considered to be a tag-jump. We tested multiple thresholds and selected 0.001% as this is when a drop in sequence reads in the negative sequencing controls occurred (Figure 8).

We summarized the noise in our dataset by plotting the total number of OTUs that were retained and those that were removed (including reason for removal; Figure 9) and did the same for the number of samples (Figure 10). After all data quality control steps described above, we plotted the total number of both sequences and OTUs before and after our data cleaning (Figure 10).

**Figures**

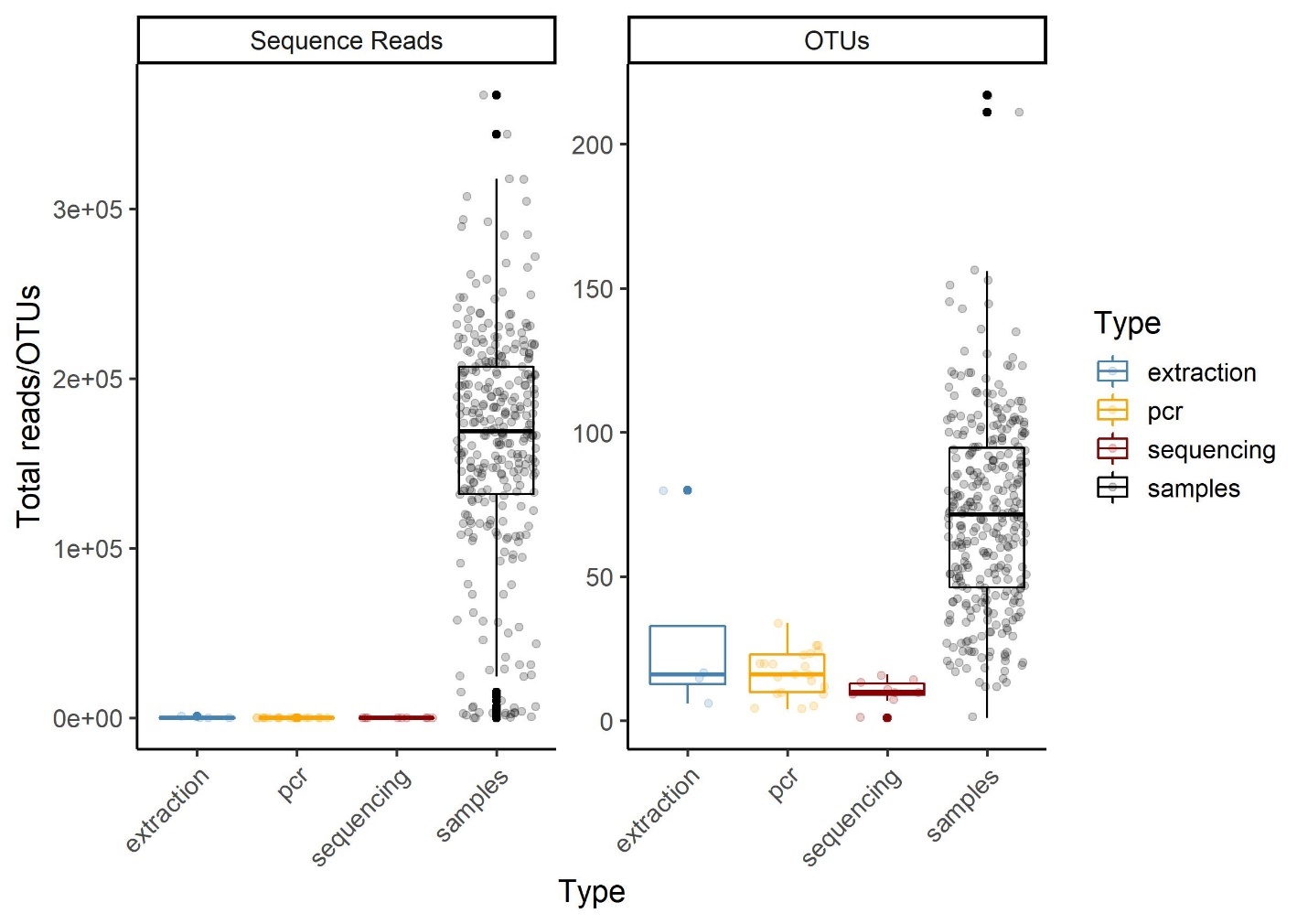

**Figure 1 –** The total number of reads and the total number of OTUs in each negative control type (extraction, PCR, sequencing) and samples to visualize the proportion of reads/OTUs in each control type.

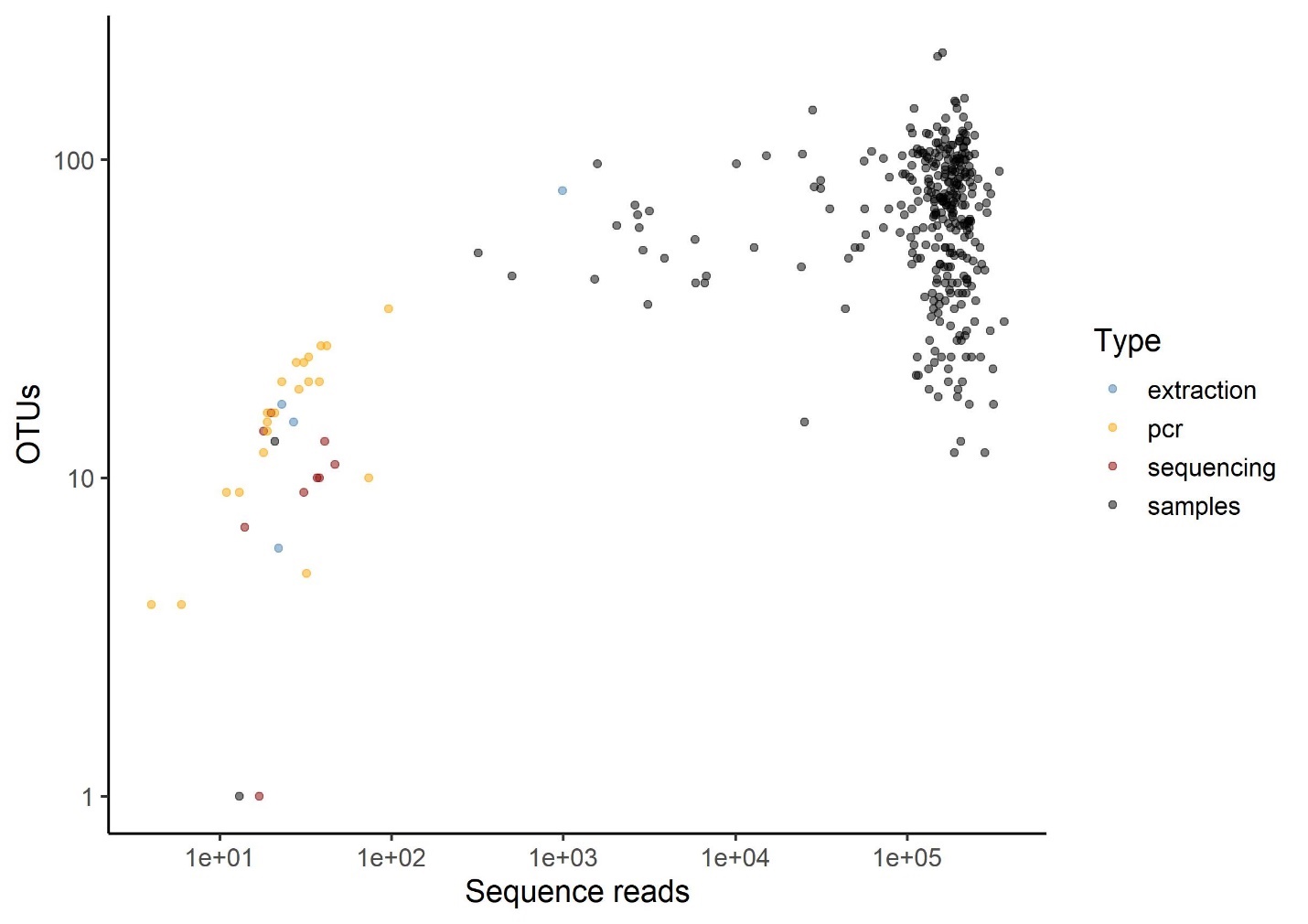

**Figure 2 –** The number of sequence reads per PCR reaction and the number of OTUs detected in that reaction. The OTUs in the samples (black circles) are not correlated with number of sequence reads and thus we can confirm that sequencing depth was sufficient.

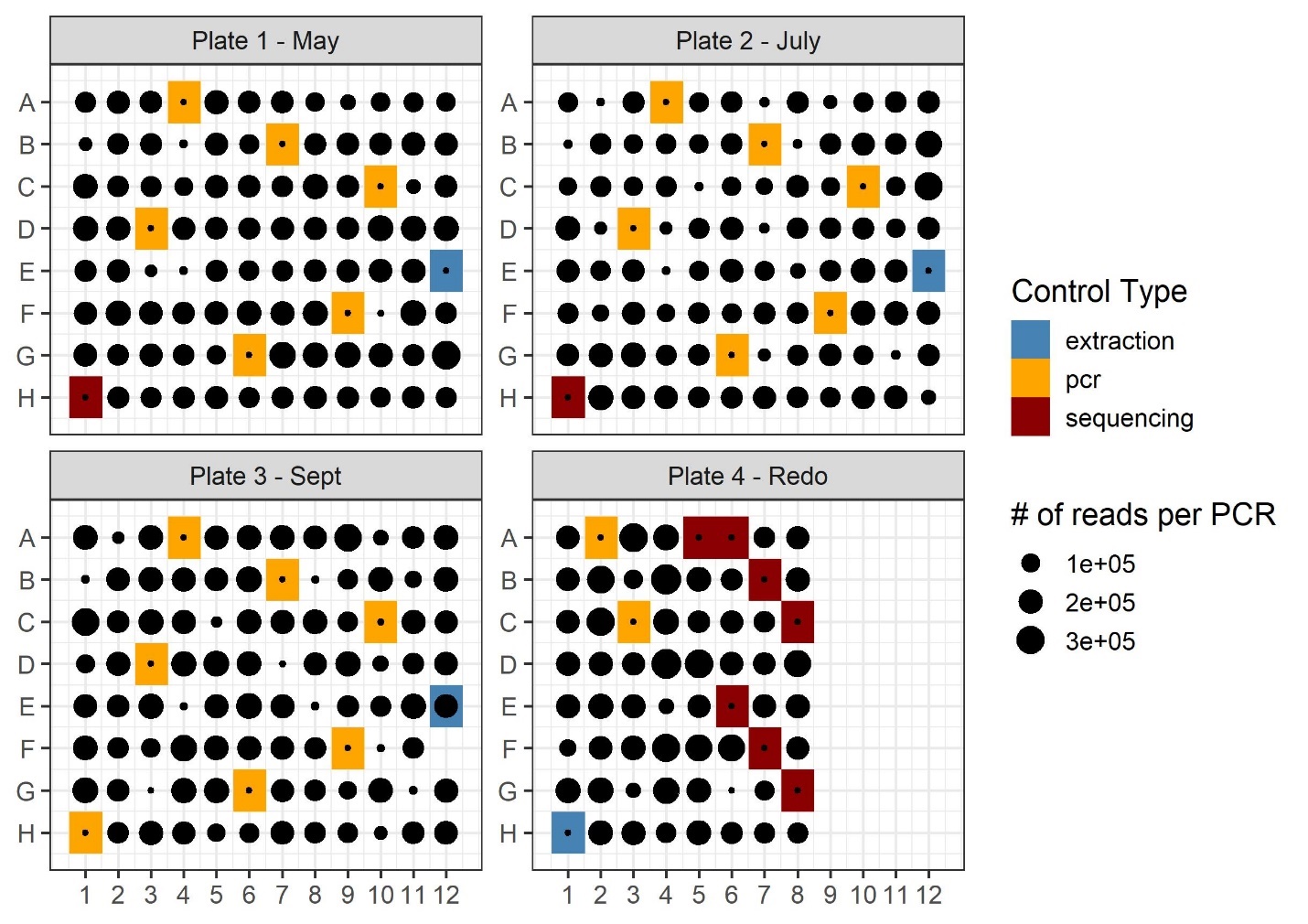

**Figure 3** – The number of reads per PCR reaction are visualized in plate format for samples and controls to confirm that there was no pattern in samples containing low sequence reads.

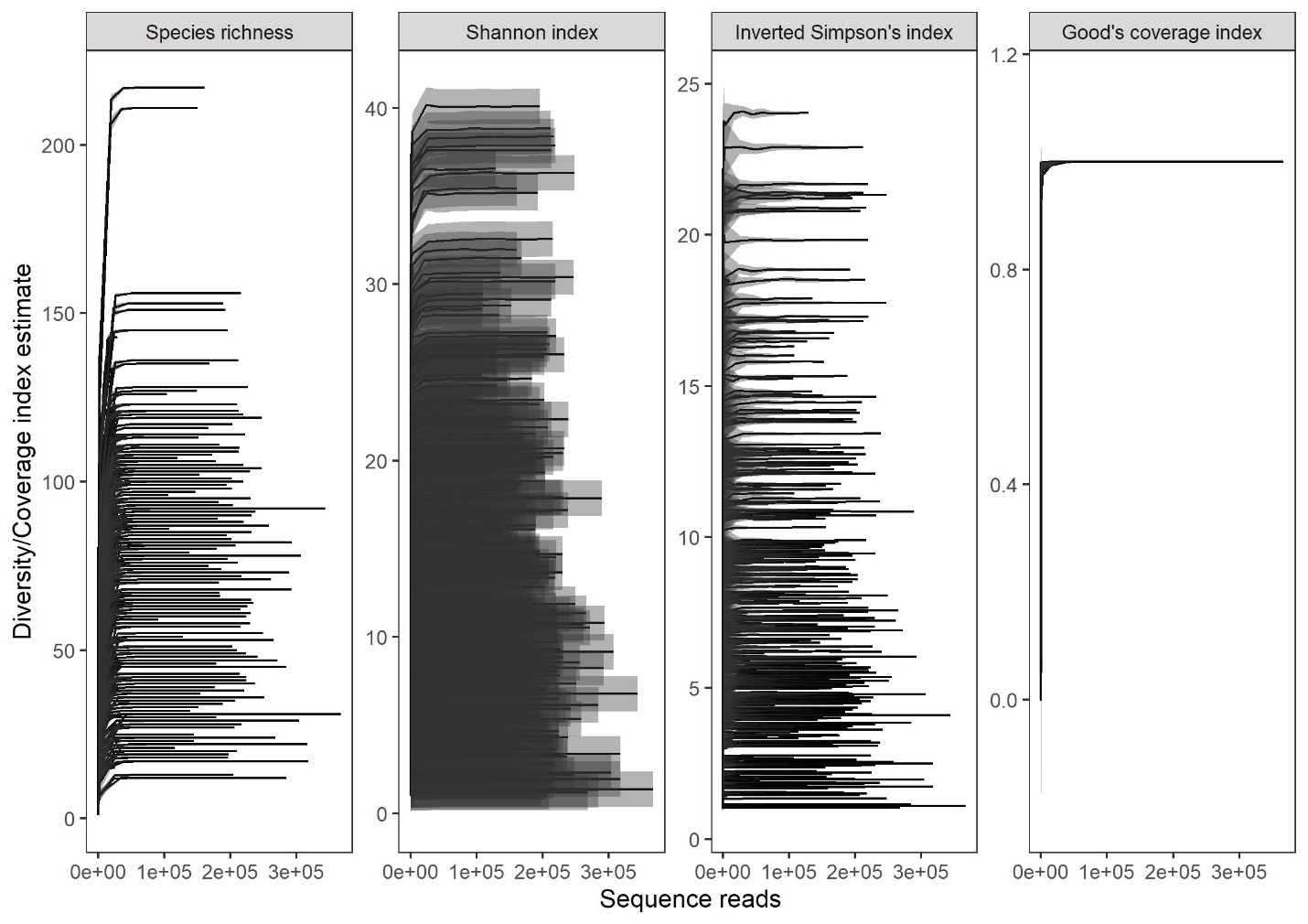

**Plot 4 –** The rarefaction curves for species richness, Shannon index, inverted Simpson’s index and Good’s coverage index for every PCR. All curves level off, suggesting read depth was sufficient in our samples.

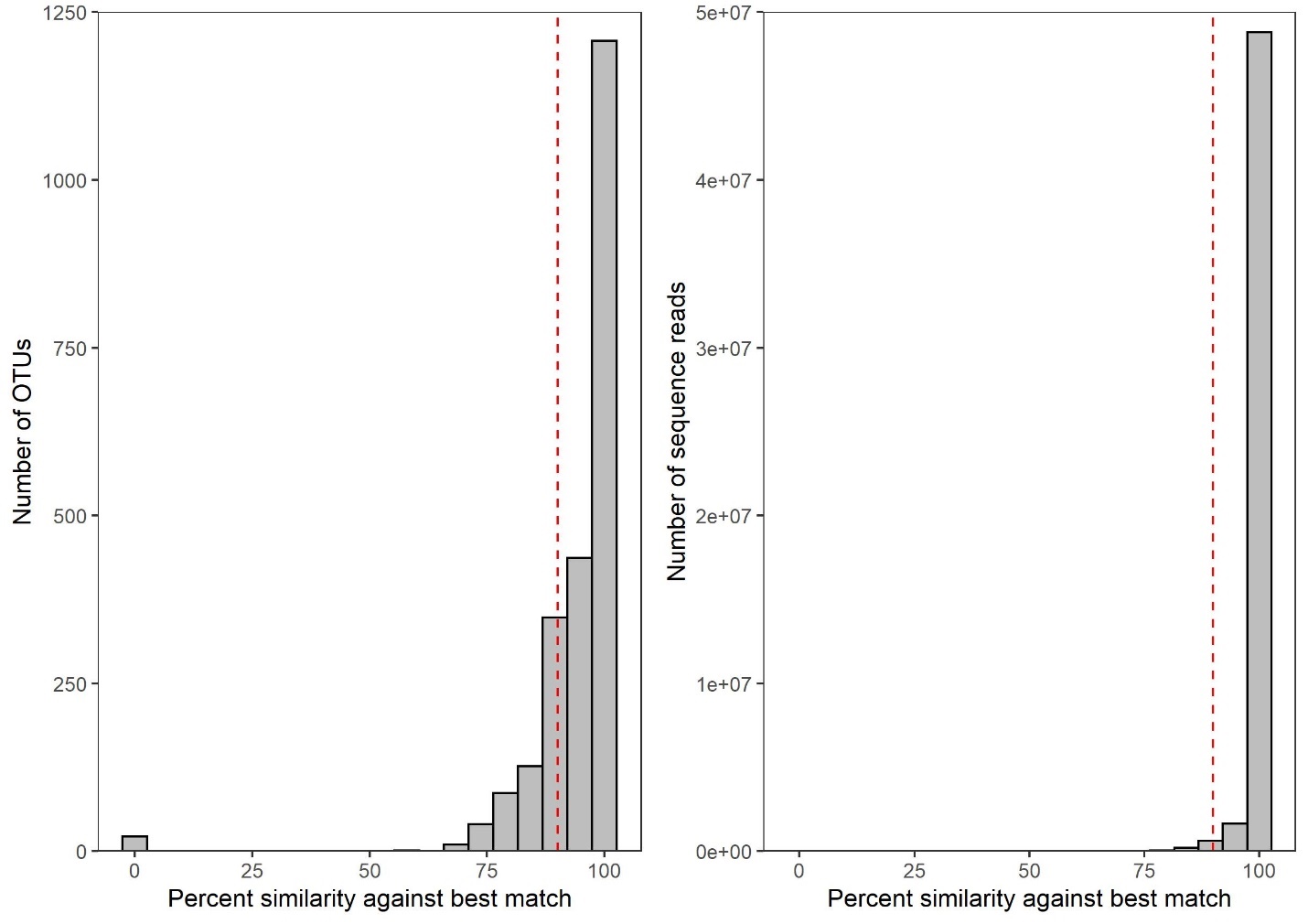

**Plot 5 –** The percent similarity of the best match against the reference database for both OTUs and sequence reads. The dashed red vertical line indicates our threshold of 90% similarity.

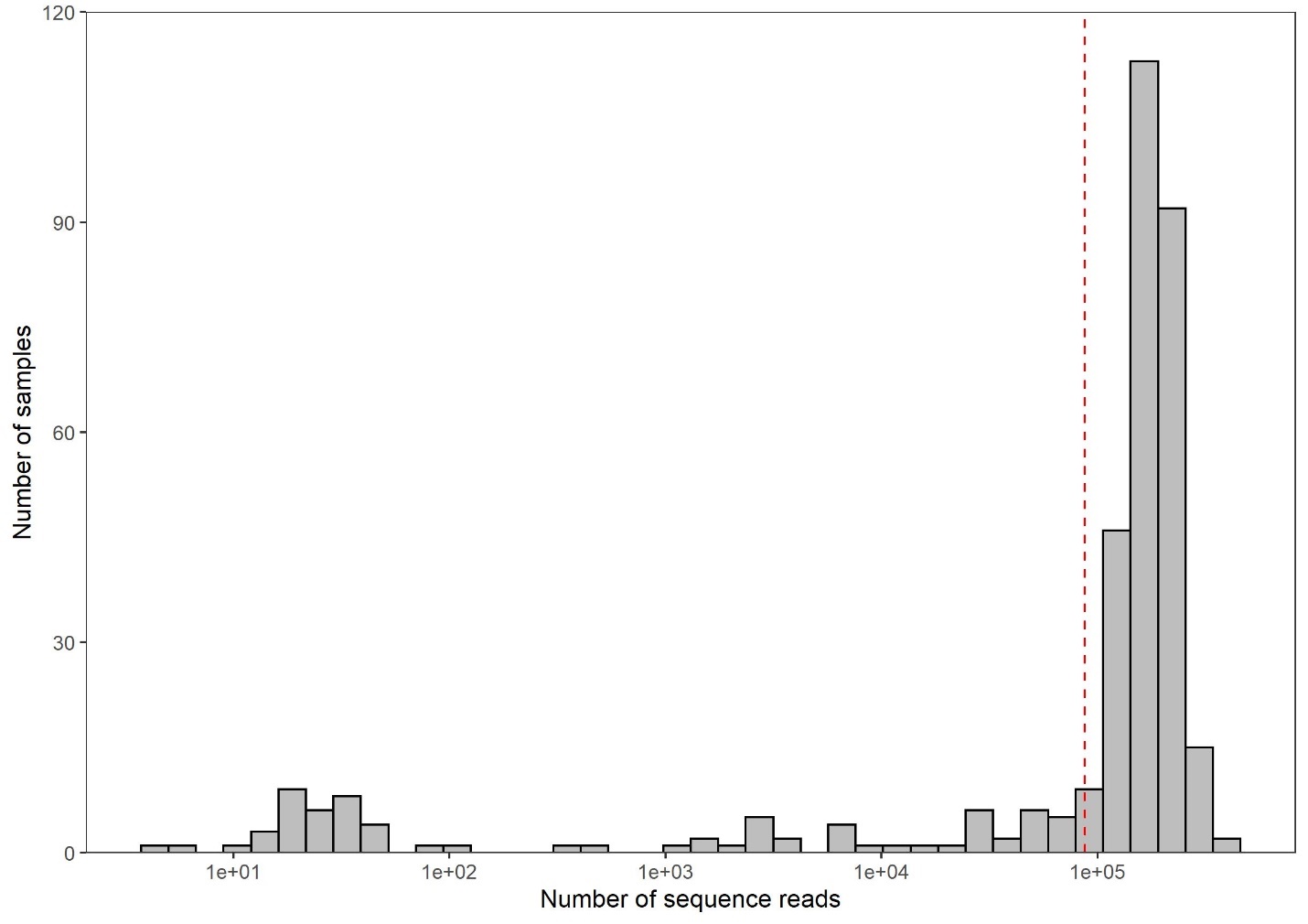

**Plot 6 –** The number of sequence reads and their frequency of occurrence (number of samples). Note that negative controls are included here. The dashed red vertical line indicates our cutoff for sequence reads (87344).

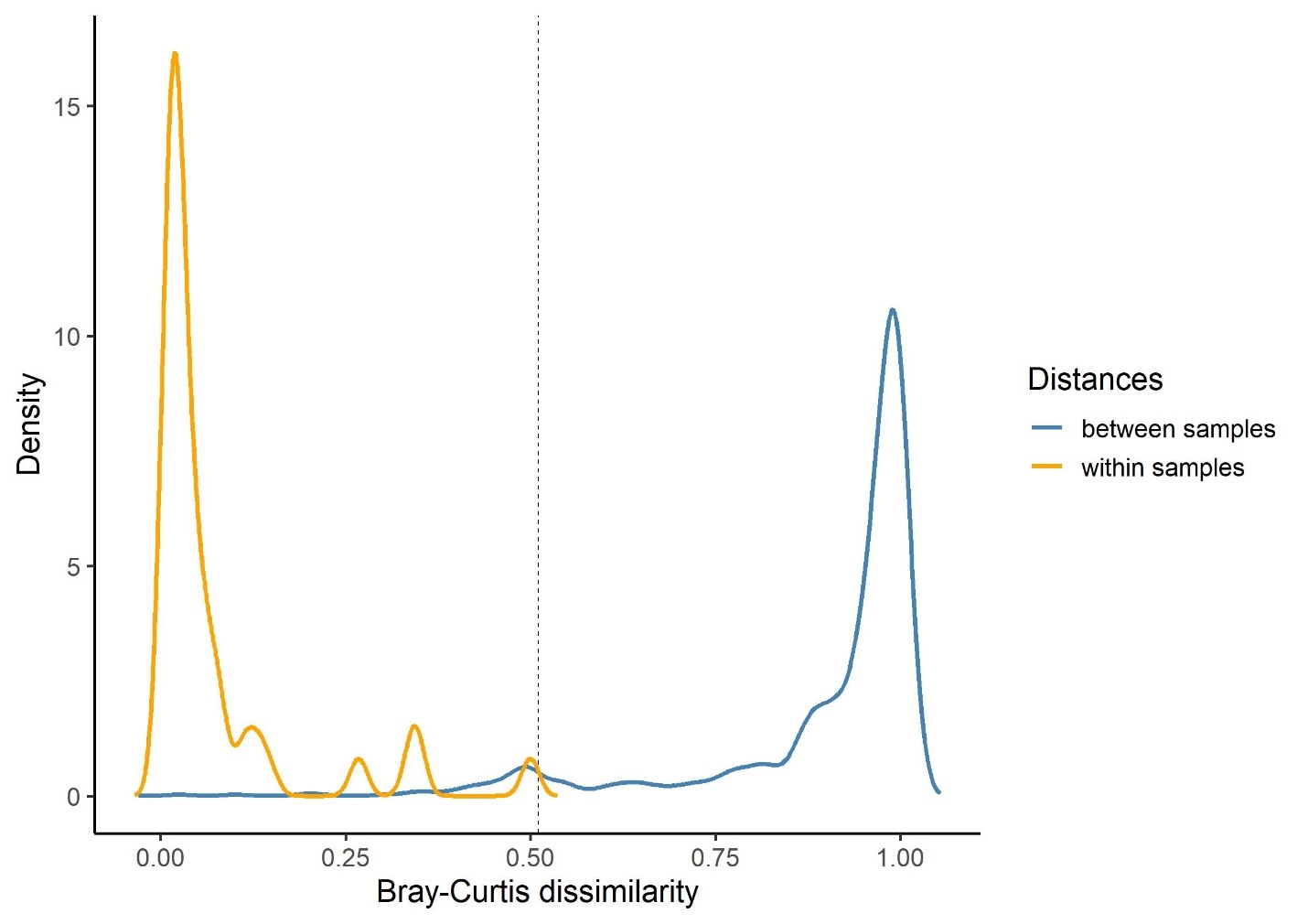

**Figure 7 –** The density of Bray-Curtis dissimilarity values within samples (e.g., PCR technical replicates) in orange and between samples (in blue). The threshold of where these values intersect (0.51) is our cutoff threshold.

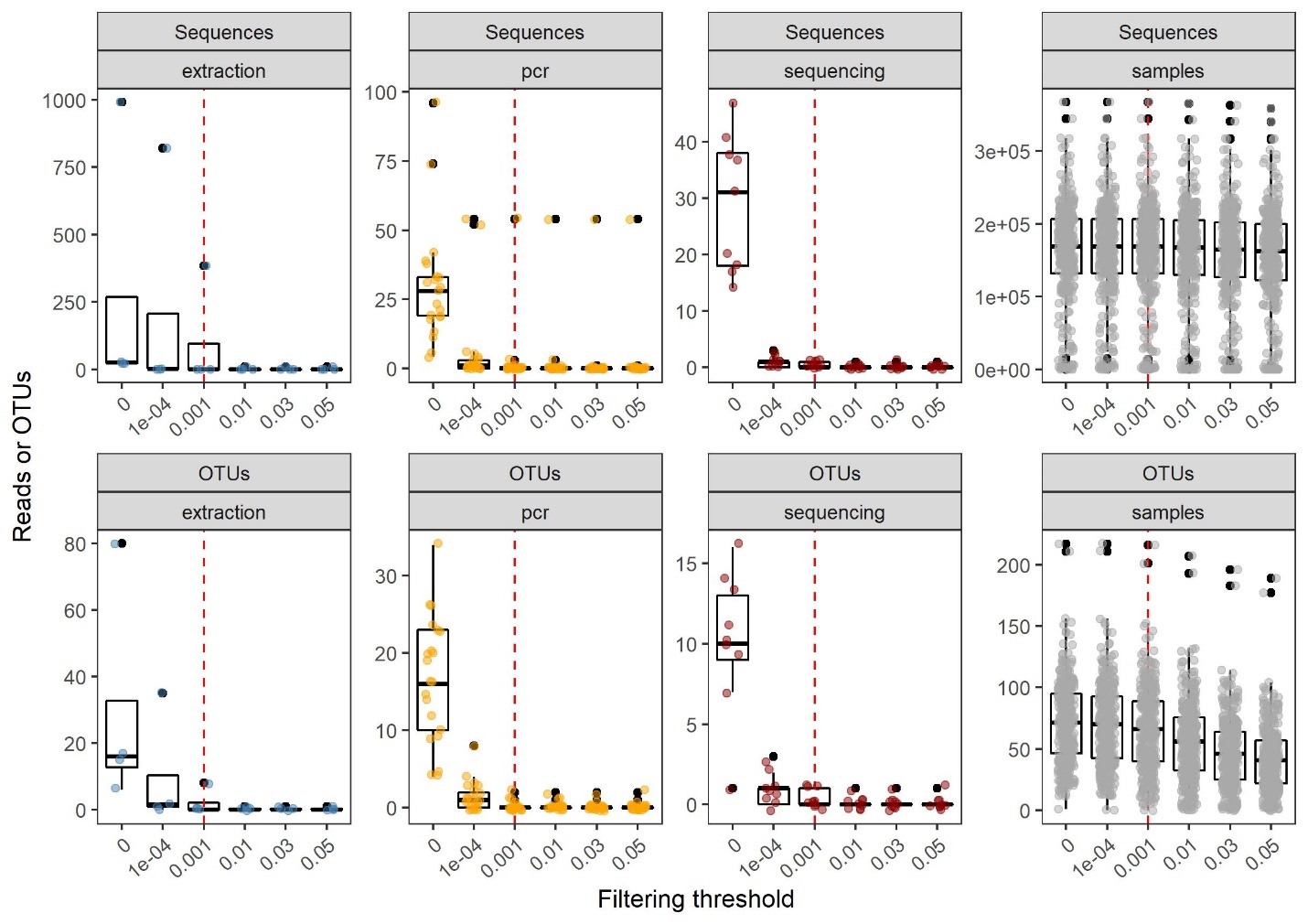

**Figure 8 –** The effects of each filtering threshold (an OTU reduced to zero abundance in a sample if its relative abundance is this percentage of the total abundance for that OTU) for both sequence reads and OTUs. The drop in reads in the sequencing controls indicate that tag jumps have been controlled for in the dataset. The red dashed vertical line represents the selected threshold of 0.001%.

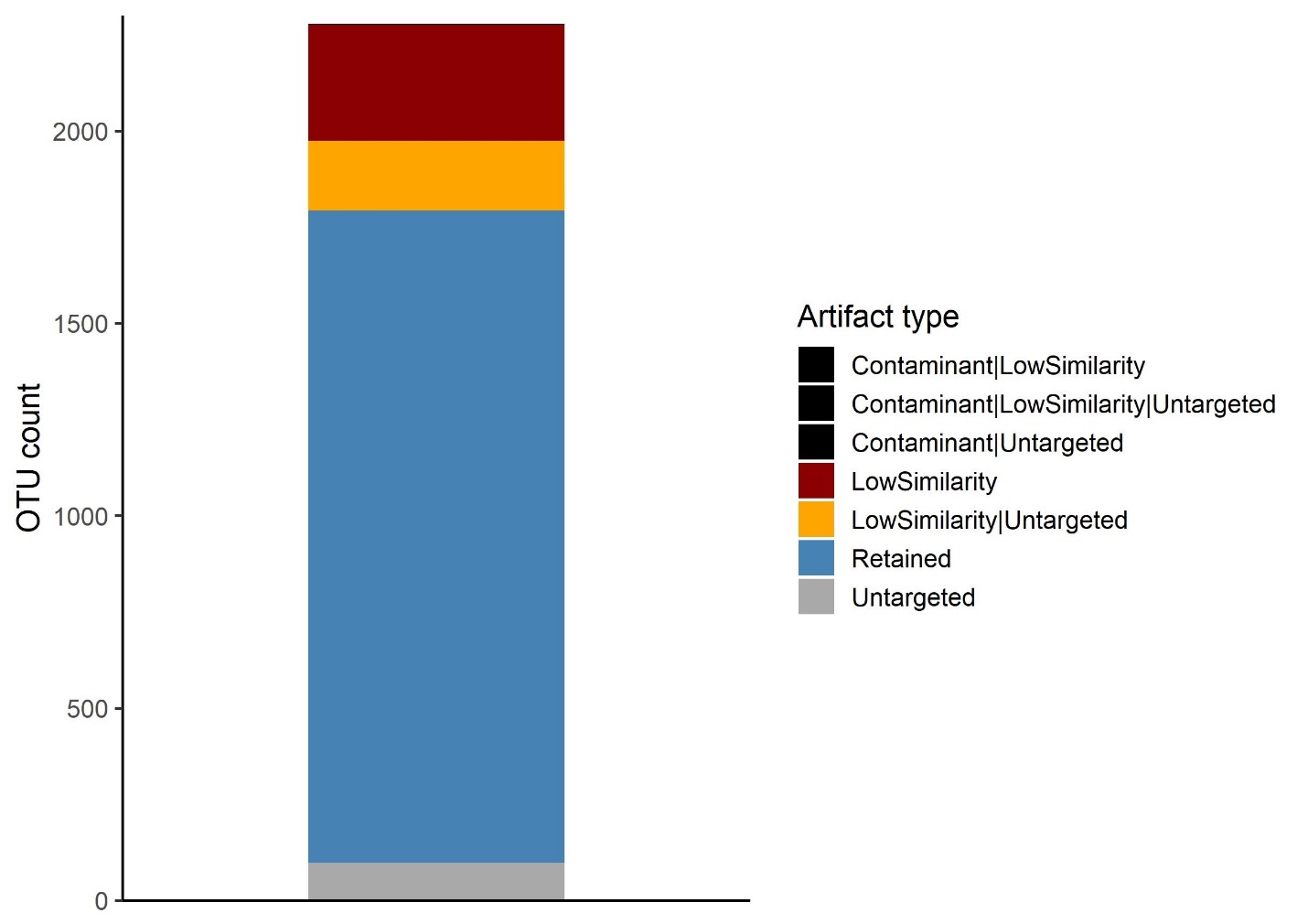

**Figure 9 –** The total number of OTUs in our dataset and the proportion that were retained. OTUs in blue were retained. OTUs in grey were removed for being untargeted taxa, those in orange were both untargeted and low similarity to the reference database (< 90%) and those in red were low similarity. Three OTUs in black are not visible on the plot, but were removed for being both contaminants and low similarity/untargeted sequences.

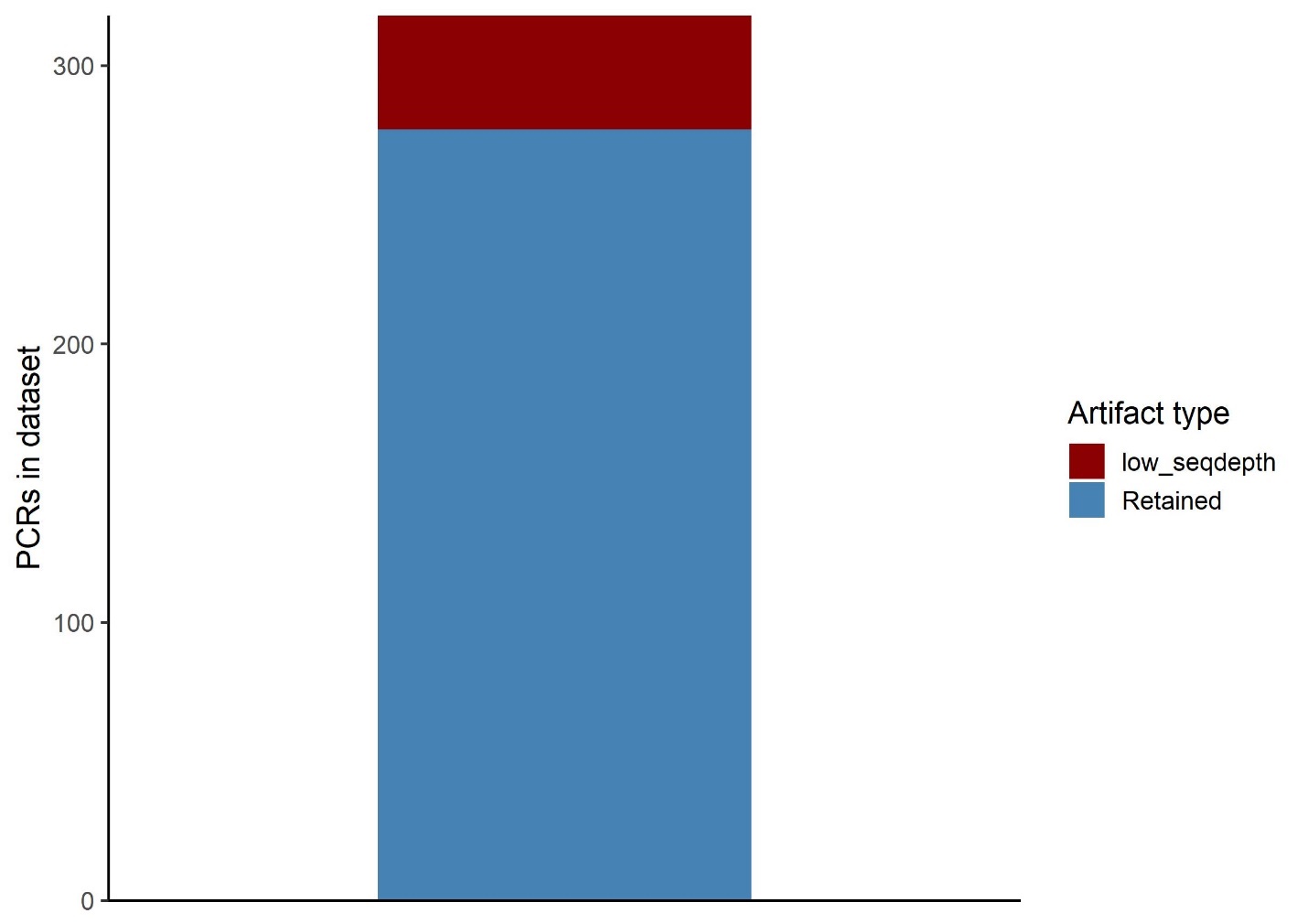
**Figure 10 –** The total number of PCR reactions (including all samples and technical replicates) in our dataset and the proportion that were retained. Samples in blue were retained and those in red were filtered out due to low sequencing depth. No samples were filtered out for other reasons checked for in our pipeline (e.g., low reproducibility, contamination issues).

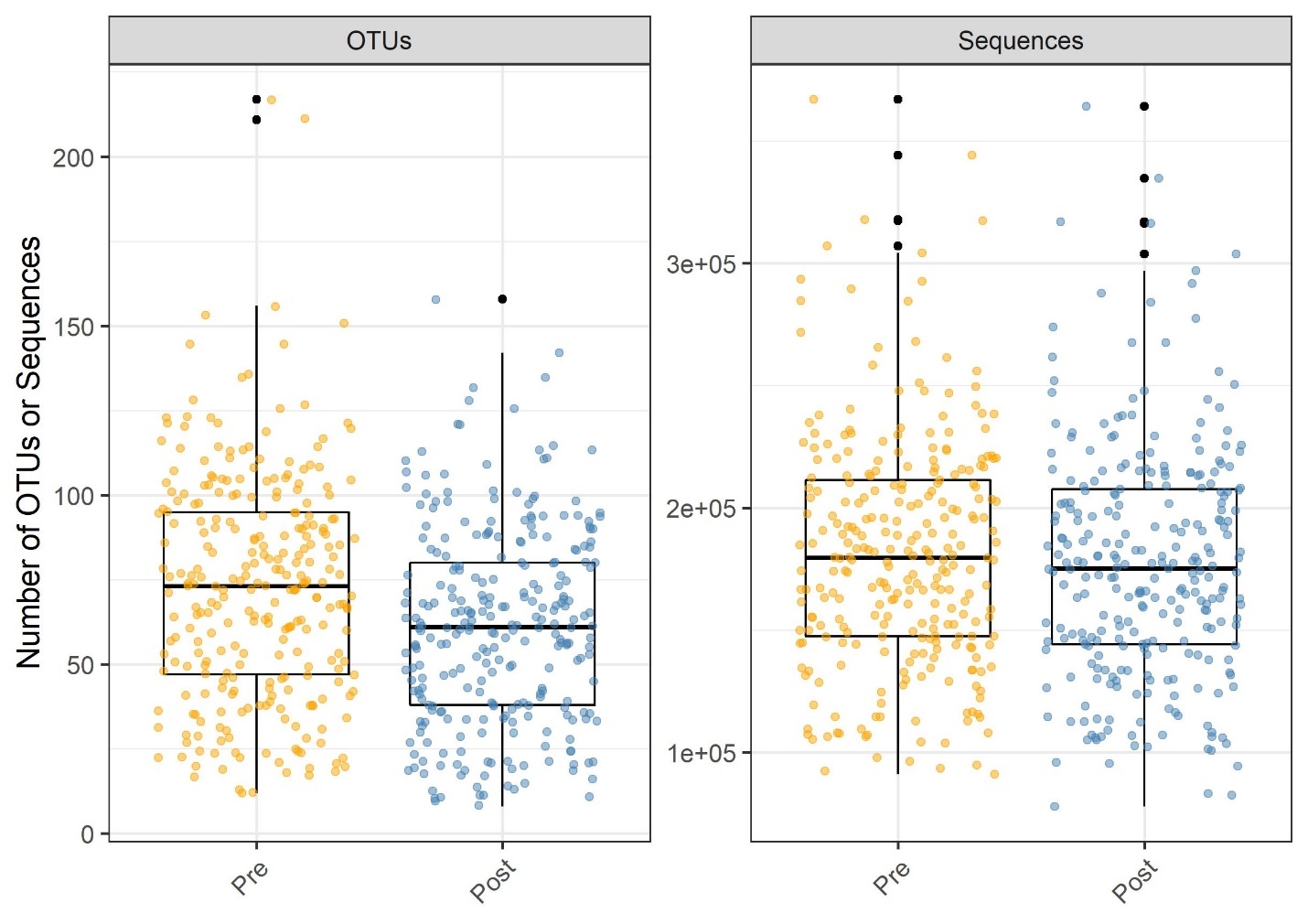

**Figure 11 –** The total number of OTUs (left) and sequences (right) in the raw data output after the JAMP pipeline (orange – pre data cleaning) and in our final dataset (blue – post data cleaning).

**References**

Chao, A., Chiu, C.-H., Jost, L., 2014. Unifying species diversity, phylogenetic diversity, functional diversity, and related similarity and differentiation measures through Hill Numbers. Annu. Rev. Ecol. Evol. Syst. 45, 297–324. https://doi.org/10.1146/annurev-ecolsys-120213-091540

Emilson, C.E., Thompson, D.G., Venier, L.A., Porter, T.M., Swystun, T., Chartrand, D., Capell, S., Hajibabaei, M., 2017. DNA metabarcoding and morphological macroinvertebrate metrics reveal the same changes in boreal watersheds across an environmental gradient. Sci. Rep. 7, 1–12. https://doi.org/10.1038/s41598-017-13157-x

Kuntke, F., de Jonge, N., Hesselsøe, M., Lund Nielsen, J., 2020. Stream water quality assessment by metabarcoding of invertebrates. Ecol. Indic. 111, 105982. https://doi.org/10.1016/j.ecolind.2019.105982

Ratnasingham, S., Hebert, P.D.N., 2007. BOLD: The Barcode of Life Data System (www.barcodinglife.org). Mol. Ecol. Notes 7, 355–364. https://doi.org/10.1111/j.1471-8286.2006.01678.x

Steinke, D., Braukmann, T.W., Manerus, L., Woodhouse, A., Elbrecht, V., 2021. Effects of Malaise trap spacing on species richness and composition of terrestrial arthropod bulk samples. Metabarcoding and Metagenomics 5, 43–50. https://doi.org/10.3897/mbmg.5.59201

Wickham, H., 2020. ggplot2: Elegant Graphics for Data Analysis. Springer-Verlag New York. ISBN 978-3-319-24277-4, https://ggplot2.tidyverse.org

Zinger, L., Lionnet, C., Benoiston, A.S., Donald, J., Mercier, C., Boyer, F., 2021. metabaR: An r package for the evaluation and improvement of DNA metabarcoding data quality. Methods Ecol. Evol. 2021, 1–7. https://doi.org/10.1111/2041-210X.13552
